# Network analysis of mesoscale mouse brain structural connectome yields modular structure that aligns with anatomical regions and sensory pathways

**DOI:** 10.1101/755041

**Authors:** Bernard A. Pailthorpe

## Abstract

The Allen mesoscale mouse brain structural connectome is analysed using standard network methods combined with 3D visualizations. The full region-to-region connectivity data is used, with a focus on the strongest structural links. The spatial embedding of links and time evolution of signalling is incorporated, with two-step links included. Modular decomposition using the Infomap method produces 8 network modules that correspond approximately to major brain anatomical regions and system functions. These modules align with the anterior and posterior primary sensory systems and association areas. 3D visualization of network links is facilitated by using a set of simplified schematic coordinates that reduces visual complexity. Selection of key nodes and links, such as sensory pathways and cortical association areas together reveal structural features of the mouse structural connectome consistent with biological functions in the sensory-motor systems, and selective roles of the anterior and posterior cortical association areas of the mouse brain. Time progression of signals along sensory pathways reveals that close links are to local cortical association areas and cross modal, while longer links provide anterior-posterior coordination and inputs to non cortical regions. The fabric of weaker links generally are longer range with some having brain-wide reach. Cortical gradients are evident along sensory pathways within the structural network.

**Author’s Summary:** Network models incorporating spatial embedding and signalling delays are used to investigate the mouse structural connectome. Network models that include time respecting paths are used to trace signaling pathways and reveal separate roles of shorter vs. longer links. Here computational methods work like experimental probes to uncover biologically relevant features. I use the Infomap method, which follows random walks on the network, to decompose the directed, weighted network into 8 modules that align with classical brain anatomical regions and system functions. Primary sensory pathways and cortical association areas are separated into individual modules. Strong, short range links form the sensory-motor paths while weaker links spread brain-wide, possibly coordinating many regions.

## Introduction

Major projects to map brain structure in a variety of animals are producing significant datasets laying out structural and functional connectivity [1] patterns that present challenges for analysis [2, 3]. Methods from network science [4, 5] can shed light on important hubs, links and modular sub-divisions. These can be characterised using tools derived from the mathematical apparatus of graph theory [2–6]. Ultimately functional circuits may be resolved, as are being painstakingly reconstructed using modern experimental probes (e.g. see [8]). The challenge is to find appropriate network models that can realistically reproduced/predict brain structural and functional connectivity data.

Here the Allen Institute for Brain Science (AIBS) dataset of a mesoscale structural connectome for the mouse brain [9] is studied using a combination of data analytical, network and visualization methods. While the data is brain wide it is at the meoscale (resolution of 0.1 mm voxels and 0.35 mm coronal plane imaging), in which links involving many neurons are probed. Connectivity data at finer scales, at the level of individual neurons and synapses, are available for smaller samples, such as with mouse retina [10, 11]. Higher resolution data also is emerging at the level of cortical layers [12]. These studies shed light on the correlation between structural and functional connections between nodes in the mouse brain. The network nodes are located at known 3D coordinates from the Allen Atlas [13–14]. Typically the sources and targets of the probed connections are classical anatomical regions that, given sampling protocols, vary significantly in size and functional significance. That non-uniform sampling presents challenges for analysis [15]. Already some network methods have been applied to the AIBS mouse connectome to find the basic network features [9, 16] or to infer hubs via their transcriptional signatures [17], especially in the cortex. These studies show consistency with a Watts-Strogatz small world and rich club connectivity. Sampling statistics, data thresholds and possible roles of weak links have also been studied [18]. While simple network theory has provided some descriptions of the mouse brain connectome it has limited ability to reveal further insights unless the network measures can capture the relevant underlying processes, such as time dependent signal flows on a spatially embedded network. It is the combination of data analytics, network analysis, 3D visualisation and process modelling that can reveal biologically relevant insights, as laid out and tested on the mouse retina connectivity data [11].

It is becoming increasingly evident that brain connectivity comprises more than a simple network. The links are directed and weighted, they can be signed: that is excitatory or inhibitory, they also can be modulated by chemicals (neuromodulators, hormones, etc) present in the extracellular space, the links have spatial extent and fit within a finite volume, and the links can be activated over time. There may be several types of networks present that operate separately or in parallel [19]: the emerging theory of multi-layer networks [20, 21] provides tools for such analysis as relevant data becomes available. In brain networks understanding is still emerging of: node and link weight definitions, information representations, signalling dynamics, and the roles of space and time in network functions.

A brain network is spatially embedded and constrained to a finite volume. It is increasingly realised that this imposes constraints [22–24] and wiring costs [25, 26] on brain networks. Meanwhile the theory of spatial networks is available [27]. The nodes are at known geometric locations that are recorded in an Atlas [13], or approximated by the experimental injection site locations [9]. Thus Euclidean distances or fibre path lengths between network nodes need to be considered along with the topological or graph path lengths (number of links or steps between nodes).

Brain networks provide pathways that transmit and actively process signals. Thus network methods that relate to information theory are likely to provide biologically relevant insights, while the topological features that underlie graph theory [28] can address other questions. Other real-world networks, such as transportation hubs or social interactions, are more suited to study by the basic topological network methods. A generic starting point for studying the stochastic aspects of signal spread is a random walk on the structural network [29]. This captures multi-step and secondary pathways and, importantly, also captures stochastic elements of the underlying processes. Beyond that better models, capturing the combined deterministic and stochastic elements, are required of the specific signalling, coding and processing mechanisms involved - as more such emerges these models can be developed and refined.

Modular decomposition methods [30] have been applied widely to brain connectivity data [1, 5] and show local clusters that are more densely inter-connected than the overall network. It has long been anticipated [31] that such modules may be hierarchically organised [32] in biological systems. It also should be borne in mind that network analysis methods commonly applied to brain data often are based on graph theory methods that rely on topological shortest paths between nodes. In a brain network that is spatially embedded, weighted and directed, and that transports and processes information, it is likely that multiple and secondary paths may contribute additional routes between nodes [33]. Random walk based methods can incorporate such multiple paths and may yield more biologically relevant modules. It also has been pointed out that plausible modular decompositions can be affected by: artefacts of discretization, threshold choice, algorithm choice, or simple ordering of nodes [15, 34]. Additionally, tree or dendrogram based methods that use correlation measures applied at a chosen cut-off level can guarantee that a hierarchy will be found.

Time dependence enters in at least two ways: the network structure may change (over longer times) and can be studies by the methods of time-dependent networks [35]; and temporal processes occur on the given network [36, 37], such as spreading phenomena or signalling. Thus not all linked nodes may be reachable within a given time frame. Just as in special relativity, where light carries signals at a constant speed, only some regions in space are reachable within a given time: there is a “space-time cone” in a brain which contains physically reachable targets [38]. Signals, carried by action potentials, spread out across the next available links in an ordered sequence of “time-respecting paths” [36, 37]. Some nodes, although linked, may not be reachable within a specified time over the available sequence of traversed links. And some network nodes may be reachable by multiple signals that happen to arrive within a small time window, so they are available for co-incidence detection and integration [39]. Other studies have used event-related networks to characterise the time evolution of brain functional connectivity [40]. Finally modular structure of the network may have some time dependence, given that links are instantiated over the signalling time-of-flight. Thus module features may evolve from an initial, local structure formed by short links to the steady state structure that includes longer, possibly brain wide, links. Separate again are representations of space and time in the brain [41], of relevance to navigation and memory: such are not considered here.

Many experimental studies of brain circuits rely on targeted experimental probes to reveal specific circuit pathways: there are many structural network links available, but not all may be functionally activated in a given situation. The disparity between functional and structural probes of brain connectivity highlights the challenge. Here, network methods can identify, or confirm, important signalling pathways in the brain such as the primary sensory routes, along with multiple secondary links to other sensory modalities and to cortical association areas. It is possible to explore the locations and timing of integration of signals from multiple senses and so explore sensory integration. Ultimately richer network models encompassing multiple network layers, signed links describing excitation and inhibition, spatial embedding, time dependence, and link activation mechanisms, may yield real theoretical insights into brain circuits and networks.

The present study integrates the methods described above to elicit biologically relevant features of the mouse brain network structure. It focuses on signal flows on the network, rather than graph methods based only on shortest paths. This paper: analyses the mesoscale mouse brain connectome, comprising weighted directed links; focuses on ipsi-lateral connections only; analyses the connectivity data by thresholding and sensitivity analysis; incorporates a spatially embedded and temporal network; employs a range network measures to identify key nodes and links; finds the network modules using the infomap method; and presents 3D layouts of network features in both the Allen Atlas coordinates and in a simplified schematic layout. It also probes the network by tracing primary sensory– motor – association signalling pathways embodied in the network, and by examining integration of signals between the association areas of the mouse cortex. More detailed network features are presented in supplementary figures, with biologically relevant results in the main text.

## Methods

### Connectivity data

The original AIBS dataset [9] was used to generate the connectivity or adjacency matrix for the mesoscale mouse brain network. The experimental data was obtained using an anterograde viral tracer, labelled with a fluorescent probe, that was injected into a source region and allowed to spread along axonal projections to linked target regions. Injection sites (source nodes) were located in the right hemisphere. Here it is the ipsi-lateral connections only that are examined. Further details of the experimental protocol are discussed in S1 Text. Given the varying sizes of sources and targets, earlier analyses have calculated a “normalized connection density” as the fluorescent signal per unit volume of the target and per unit volume of the source [9, 16, 18]. However most network analysis typically deals with all weighted links between nodes, rather than a density of links. Although some new methods allow for node weights, for example arising from city areas or brain region volumes, as well as link weights [42–43], the significance of link density is not well developed in network models, so it is unclear how to apply this to brain data. Here we use the usual network model [6, 28] that incorporates varying node sizes and total link weights. The original raw data is the normalized connection strength, originally termed w_XY_ [9], being the volume of fluorescent voxels detected in the target region arising from the volume of tracer injected in the source region, and was reported in Suppl. Table 3 of [9]. Whether this is the true network connection strength between the two regions is a question of volume sampling in the source region, as presented in S1 Text.

It is helpful for subsequent theoretical analysis to further normalise the measured values to some representative strength, so the link weights are hereafter scaled to a unit value and are dimensionless. This is consistent with the common observation that many scientifically measured quantities have a typical size or scale [44]. Additionally well normalised quantities allow for more robust and efficient computation. As noted below the maximum and minimum measured values are possible outliers, so do not form a reliable basis for normalisation; and a layered data acceptance or rejection approach may be needed for concordance between structural and functional measurements. Thus some reliable, intermediate value needs to be chosen. Following the lower limit adopted in Figure 3 of [9] it is convenient to rescale the raw data by a_0_ = 10^−3.5^ (mm^3^), which is then taken as a unit link weight. The elements of the adjacency matrix, A are then well conditioned for matrix computations. Calculated network properties are unaffected by this choice.

S1 Figure plots all the original raw data and highlights obvious outliers. A guide to thresholding the data is provided by tradeoffs between True- and False-Positives rates in comparisons between different types of connectivity measurements. Details are provided in S1 Text. Those studies [45–46] suggest that the top 40%, by weight, of links provides an optimal comparison between structural and functional measurements. Thus the present study focuses on the top 40% of structural links by applying a link weight threshold, at weight 78 using the normalisation described above and in S1 Text. The sensitivity of results to threshold choice is investigated and the weak links are studies separately.

### Link distances

The present analysis also required the distance matrix which records link lengths between nodes. Geometric distances between nodes (anatomical regions) were taken from the coordinates available online with the Allen Atlas [13–14]: the Waxholm space delineation 2012 (V2) for the C57BL/6 mouse was used, since this was then contemporary with the connectome experiments and is still used in various AIBS online products. A later, higher resolution, edition, the Allen Mouse Brain Common Coordinate Framework version 3, was used to check selected results, noting the different coordinate origin. In the present application only distances between nodes were required. The centroid (median values of x, y, z) of each region was used as a representative node coordinate; typically this would be close to the centre of mass, except for regions of unusual shape. Available 3D volumes of Nissl-stain sections and labels at voxel level, along with NIFTI tools in Matlab [47] enabled comparison between the three sources. Typically differences up to 0.2 mm were encountered, with larger, extended anatomical regions (eg. MO) being more difficult to characterise by a single coordinate. Here geometric effects are limited to the influence of link length and strength on network features – rather than geometric constraints, such as a finite brain volume.

### Network analysis and modular decomposition

The network analysis methods were described previously and tested on the mouse retina connectivity data [11] and follow standard practice [1, 5–7, 28]. The nodes of network are the 213 anatomical regions corresponding to the selected experimental source (injection sites, or combinations thereof) and target sites [9]. The resultant array of source-target link strengths forms the adjacency matrix [5, 6, 28] for subsequent analysis. Its entries, A(i,j) encode the weight and direction of links between nodes i and j, chosen from the 213 anatomical regions. The matrix is not symmetric since the network links are directed. The diagonal elements are set to zero since self-loops, present in the original data, are ignored in the present analysis. Thus there are (n−1)(n−2) = 44,732 possible directed links between the 213 chosen anatomical regions. The adjacency matrix was normalised and thresholded as described above and in S1 Text (cf. S1 Figure).

Standard network measures have been used on brain data [1, 5, 48], and the mouse data [9, 16], to rank the importance of nodes and links. The weighted in-degree and out-degree of each node (i.e. node “strength”) is the simplest such measure, being the column sum and row sum, respectively, of the weighted adjacency matrix. These quantities can be calculated directly from the published Allen data [9]. Other measures are the node betweenness centrality (nBC) [49] and edge BC (eBC) [6, 28]. They are based on graph shortest graph paths or geodesics, as distinct from the geometric path length that reflects the distance travelled between nodes along links in physical space. The random walk centrality measure (rwC), developed in other contexts [33], may be relevant to brains where signals can spread by both direct paths and also via secondary links. Analysis of complex metabolic networks has generated the concept of hub nodes [50], based on the participation coefficient, P which measures the diversity of a node’s out links to other network modules (see below). The concept has proved useful in analysis of brain networks [51]. All measures were calculated by standard methods and toolboxes using available Matlab codes [52–54].

### Modular decomposition

Modular decomposition of the mouse connectome was attempted by the classical methods: spectral partition of the Laplacian matrix and of the modularity matrix, k-means clustering of the leading eigenvectors of the adjacency matrix or its associated Laplacians [55–56], agglomerative and divisive methods [57], and the Louvain method [58] which iteratively agglomerates nodes. Most of these methods utilise network topology measures (e.g. node BC, edge BC) that reflect shortest paths on the topological network. These clustering algorithms do not always generalise to weighted, directed networks, as required here. The number of modules sought has to be specified in advance, and may be limited to 2 for some of these methods. The InfoMap algorithm [59–60], by contrast, uses information theoretic concepts (Hoffman code length) as a proxy for random walks on the network [59]. It is well suited to studying spreading phenomena on networks [61], and aligns well with an essential feature of a brain network, to host signal flows on the network of neurons and synapses. It also does not require the number of expected modules to be specified in advance - that emerges naturally from the calculation. Modules found by this method are essentially groups of nodes in which a random walker spends a larger fraction of time. This method has been shown to perform well on a range of test cases [59–60], and has been applied successfully to the mouse retina connectivity data [11], thus Infomap is the primary used herein. An efficient C++ code and online apps are available [62]. The Louvain and InfoMap algorithms generally produce similar results and have proved useful for biological systems, including brain networks. Both methods can uncover hierarchical modular structure. Newman’s Modularity metric, Q [55–56] was used to compare different partitions against a null model (e.g. random or configuration model).

### Temporal Networks

Here the analysis and visualization of temporal, or time evolving, links involved choosing a specified source node[s], finding the list of target nodes, via in- or out-links, then re-ordering these by increasing link lengths, and then drawing them in distance, or equivalently time, sequence. For links longer than ~5-6 mm (about half the brain size for the mouse) it is possible for 2-step links to be simultaneously active in the distance (time) window and thus warrant examination. The mid node was already directly linked to the source node, so credibly signals could continue on to/from a relevant target/source node. For those second links the synaptic delay needs to be included to correctly describe signal traversal across two links. Signal delays over axons and synapses are discussed in S1 Text, with 1 mm/millisec (mm/ms) being chosen as a typical velocity. A representative synaptic delay of 2 ms adds an equivalent path length of 2 mm for the purposes of tracing 2-step links.

### Schematic coordinates and 3D visualization

Network plots usually have too much visual complexity (cf. S4 Figure), so a simplified 3D schematic layout of the mouse brain network was sought. A set of schematic coordinates was developed, in which the original 213 anatomical regions, located at the Allen Atlas coordinates, were condensed onto 55 locations that each aggregated related and nearby regions, as detailed in S1 Text. A benefit of this approach is that related links, between nearby and related nodes, are bundled into a few pathways, enabling easier navigation of the 3D visualizations. Surface colours match those presented in the Allen Brain Atlas sagittal slice images [63]. All figures were produced in Matlab, with all codes available [64].

### Strategy

Here four strategies are adopted: network nodes are displayed at simplified schematic coordinates (cf. S1 Text, S3 Data and S10 Figure), which serves to reduce the number of link crossings and so reduce visual complexity; a number of landmark nodes were examined (e.g. high In- or Out-degree, high flow, or hub nodes); and links are selected from either the strongest (e.g. top 40%) or weakest present; next functionally relevant pathways are examined, such as the primary sensory paths, links with the cortical association areas; or links with the motor regions. The latter approaches are informed by the many experimental probes of the sensory and association cortex pathways. These, historically, have produced basic insights into nervous system and brain function [65]. Similarly network methods allow computational probes of links with key sensory and cortical regions.

## Results

### Connectivity strength versus distance

The links weights used here are the original normalized projection strength [9] (cf. S1 Figure and S1 Text) suitably normalised to a unit value (cf. Methods). Figure 1 displays the link weight vs. distance data (cf. Methods) as a log-linear plot, omitting the 10 extremely low points in S1 Figure, and with the link density normalised by a nominal threshold value of 10^−3.5^ (mm^−3^) [9]. A linear fit to the plot shows the distance trend and the significant dispersion in the data. Using a kernel regression model to infer voxel level connectivity [66] also found a linear trend for the whole brain ipsi-lateral data, with a log(weight) - log(distance) slope of −3.16, and residuals best fit by a log-normal distribution. Note that the comparable log-log fit to the data in Figure 1 yields a slope of −2.42, with R^2^=0.18 and 95% confidence interval of [−2.49, −2.34]. The trend is reminiscent of that found with diffusion MRI of human brain [67]. In that study the overall trend was quadratic, spanning 200 mm, however at the scale of the mouse brain (~12 mm) it was essentially linear, with any quadratic term making a negligible (<2%) contribution. De-trending the data, by subtracting out the linear fit, shows that the residual log-weights appear to be skew-normally distributed (Matlab dfittool) about the local mean values (now zero at each distance), with overall standard deviation 1.31. Longer links (>5mm) show slightly more dispersion (standard deviation of 1.4, vs. 1.13) than shorter links (<5mm). This is in contrast to the larger difference noted for human brain over the much longer distances, where the dispersion of log-weights increased with link distance.

**Figure 1.**
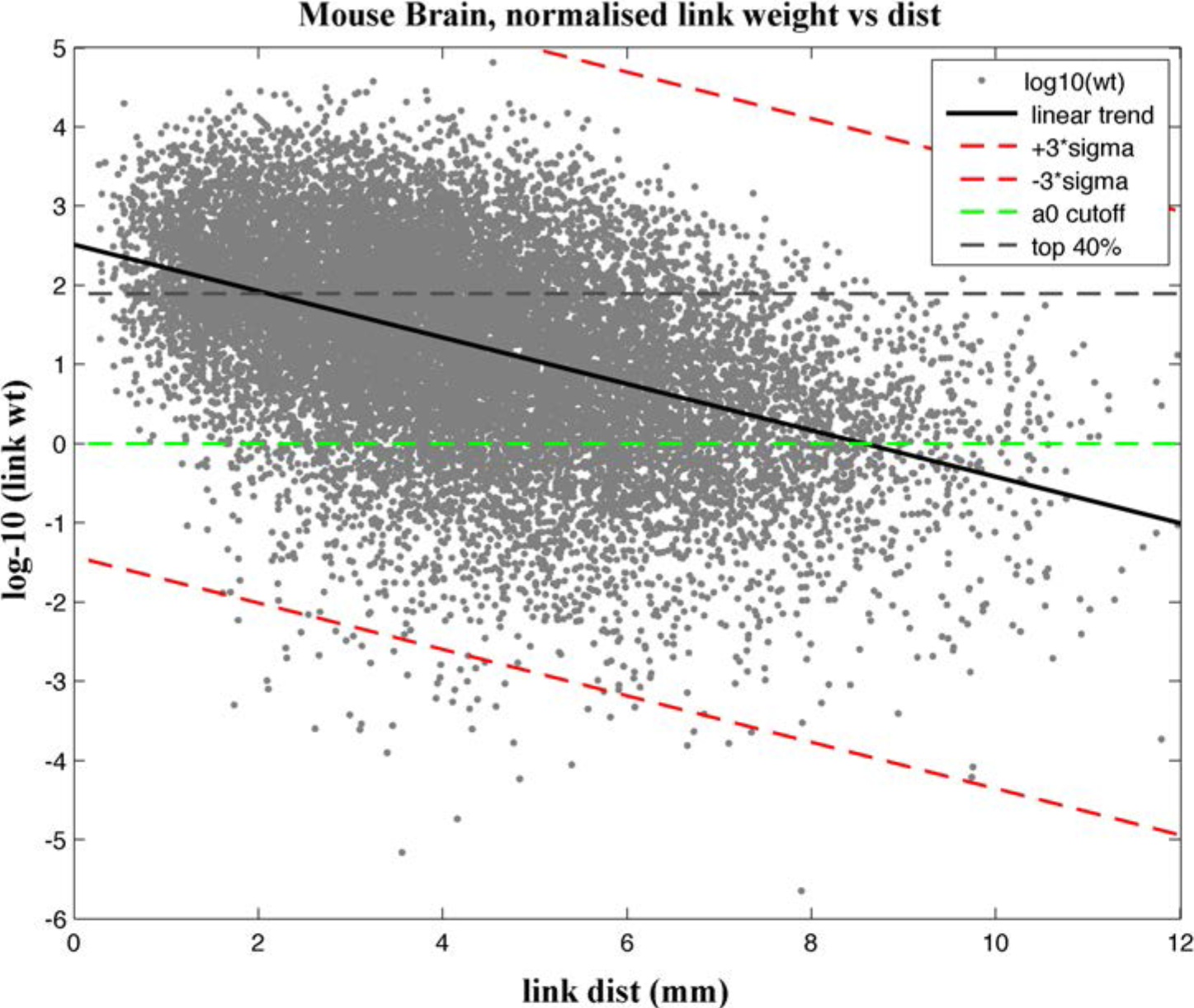
Link weights vs. distance. Normalised link weights, plotted as log_10_(weight) vs. link distance; and log-linear fit trend (black solid line) fit by the equation log_10_(weight) = 2.509 −0.293*dist(mm) with R^2^=0.20; 95% confidence interval for the slope is [−0.307, −0.289]. Also shown are: the 3-sigma lines (red, dashed), the unit weight (i.e. log_10_(weight) = 0, green dashed line) and the top 40% (gray dashed line) cut-off values.

Figure 1 also shows the local data mean (the linear trend line) and the 3-sigma trend lines (cf. S1 Text for the correct sigma of log transformed data). Often in data analysis points lying beyond the 3 standard deviations are omitted as outliers. Here 46 data points lie below the lower 3-sigma line so might be omitted as statistical outliers. Note this criterion now depends on link distance, so some care is required in eliminating weak connections. It is now clear that most of the data lies within the 3-sigma curves. Note that the strongest In-link, to CP (Caudoputamen), which appears to be an outlier, does lie within the local 3-sigma boundary. This node is also by far the largest sampled anatomical region. That makes it problematic to normalise the links to the strongest weight so that the normalised weights vary between 0 and 1, as is often used. Figure 1 plots two credible link cut-off strengths: at weight 1 (corresponding to Oh’s [9] normalised projection strength of 10^−3.5^ mm^−3^), and at the 40% link weight cutoff (cf. Methods and S1 Text). The plot makes it evident that a simple, uniform threshold discards data in a skewed manner, with shorter stronger links preferentially included, and more of the longer weaker links discarded.

### Adjacency matrix and networks characteristics

The raw connectivity, or adjacency, matrix normalised as described above and omitting the 90 self-loops contained 16,864 directed links between the 213 separate regions: 38% of all possible links. The initial analysis [9] utilises an effective cut-off strength of 10^−3.5^ mm^3^ (approximately 1/3 of a voxel). The weakest remaining link was assigned weight 1, so the strongest link has weight 64.5k, spanning 5 orders of magnitude as reported previously [9, 16]. The significance and possible roles of the weaker putative links is examined separately below. The adjacency matrix is listed in S1 Data.

S2a Figure shows the distribution of the weighted in- and out-degree (node strength) of the 213 anatomical regions (network nodes). The distribution appears heavy tailed, but with only 15 nodes having a weighted degree (in or out) greater than 10,000. In that case the mode (middle value) is a more robust estimator of representative links weights than the mean, which is unduly influenced by the larger values. The link weights have mode values of 264 (in) and 79 (out), both significantly below the strongest link weights. One node (#36, CP) has a very large weighted in-degree (64k) and might be considered a statistical outlier, being near 3 standard deviations above the local mean. It represent a quite large anatomical region, comprising 24k voxels with an indicated geometric extend of approximately 3 mm. It also is a significant outlier in a plot of wt- in-degree vs. number of in-links (not shown). The unweighted node degree (binarised adjacency matrix), that counts the number of links regardless of weight, is presented in S2b Figure. The mean number of both in- and out-links is 80, with nodes having between 3 and 180 Out-links, and between 50 and 120 In-links. Interestingly both link count distributions appear to be nearly Gaussian, with the out-distribution more skewed to the right. This suggests the speculation, in light of the Central Limit Theorem, that links might originate from a process having multiple random features, with link weights subsequently developing by more specific biological mechanisms. At mesoscale resolution, with 213 nodes, there are not enough data points to distinguish if the degree distribution follows an exponential or power law.

S3a Figure shows the distribution of graph shortest paths for the full network. These are the shortest path length (number of links traversed) between any two nodes. In a graph theoretic sense the network is compact: approximately one third (31%) of node pairs are separated by one link, with the balance (68%) reachable by two links; a very small number (232 pairs) require 3 links. Any node can be reached from any other in at most 3 links. S3b Figure shows the distribution of link lengths in geometric space: it appears to be a normal distribution, skewed to the right. The peak shows the most probable link length is short ranged (~3-4 mm), while the long tail to the right shows a still significant number of very long links (~10 mm), almost spanning the full brain.

Key nodes are listed in S1 Table, with the top 20 nodes ranked by probability flow, along with other network measures (nBC, eBC, rwBC, P). These nodes are most frequently visited by a random walk on the network. Many have high degree and number of linked (i.e. un-weighted degree). Some, but not all, have high nBC. S2 Data lists all nodes with detailed information and network measures. Potential hub nodes [50, 51] were identified by the participation coefficient, P which measures the diversity of a node’s out links to other network modules. More than half of the mouse brain nodes (113 out of 213) have P > 0.5 (within a range 0 - 1), indicating some hub-like character. Of these, five nodes also have very high weighted In-degree and random walk flows; while six nodes have very high weighted Out-degree. Together these measures may help to identify connector hubs. Likely hub nodes are listed in S2 Table. Such hubs often are evident in the Figures. Amongst the high participation coefficient nodes are 21 cortical regions, including most of the association areas, the visual and auditory cortices, and the secondary motor region. Three way comparisons of probability flow and participation coefficient to both In- and Out-Degree (figures not shown; cf. S2 Table and S2 Data), are informative. A few nodes (MRN, PAG, SCm, LHA, ACB) stand out by having high flow, with many others clustered near low flow and degree. Participation coefficient highlights other nodes (eg. CLA, CLI, PP, GPi, MGd, MOs, ENTl) and tends to spread nodes out more evenly. In each case CP is possibly an outlier due to it being the largest sampled anatomical region. A similar comparison using node BC is less informative since more nodes are clustered near zero. Overall it appears that these standard network measures, applied at this spatial resolution, provide only a partial guide to functional significance.

### Modules

The results presented below all use the InfoMap method applied to all 14,103 links of weight 1 or larger (cf. Figure 1). Ranking of nodes in order of probability flow correlates well with random walk centrality and weighted in-degree (cf. S1, S2 Tables). In part that reflects the algorithm design. However such ranking does not correlate well with shortest paths node BC, with the latter likely dominated by numerous long-range weak links. The Infomap calculations were robust to the choice of filtering (cut-off) of link weights: cut-off weights of 1, 78 (top 40% of links), 0.1 and 0.01 produced either identical results (for 1, 78) or only minor differences (for 0.1, 0.01), suggesting that the modular decomposition is robust. In all cases the major modules and their leading (i.e. highest flow) nodes were identical.

Figure 2 shows a comparison of the mouse brain anatomical regions and the calculated network modules. The regions are represented by the 3D convex hull surface that encloses all member nodes, plotted in the Allen Atlas coordinates. Note that the convex hull encloses the node coordinates that are the centroid coordinates of the anatomical region, rather than covering the full extent of those regions. Thus the region-enclosing surfaces are not space filling: this improves the visibility of the 3D graphic. Surface colours match those presented in the Allen Brain Atlas sagittal slice images [14]. The modules are presented in 2D only to reduce confusion in the image. Modules are represented by the 2D slice of the convex hull (enclosing all module members) drawn at a height (in the Dorsal-Ventral axis) corresponding to the centre of mass (CM) of the module members. Note that there is significant, but incomplete, correspondence between modules and the major anatomical regions. Overall 74% of all links are between modules, with mean length 4.3 mm, while 26% are within modules, with mean length 2.8 mm.

**Figure 2.**
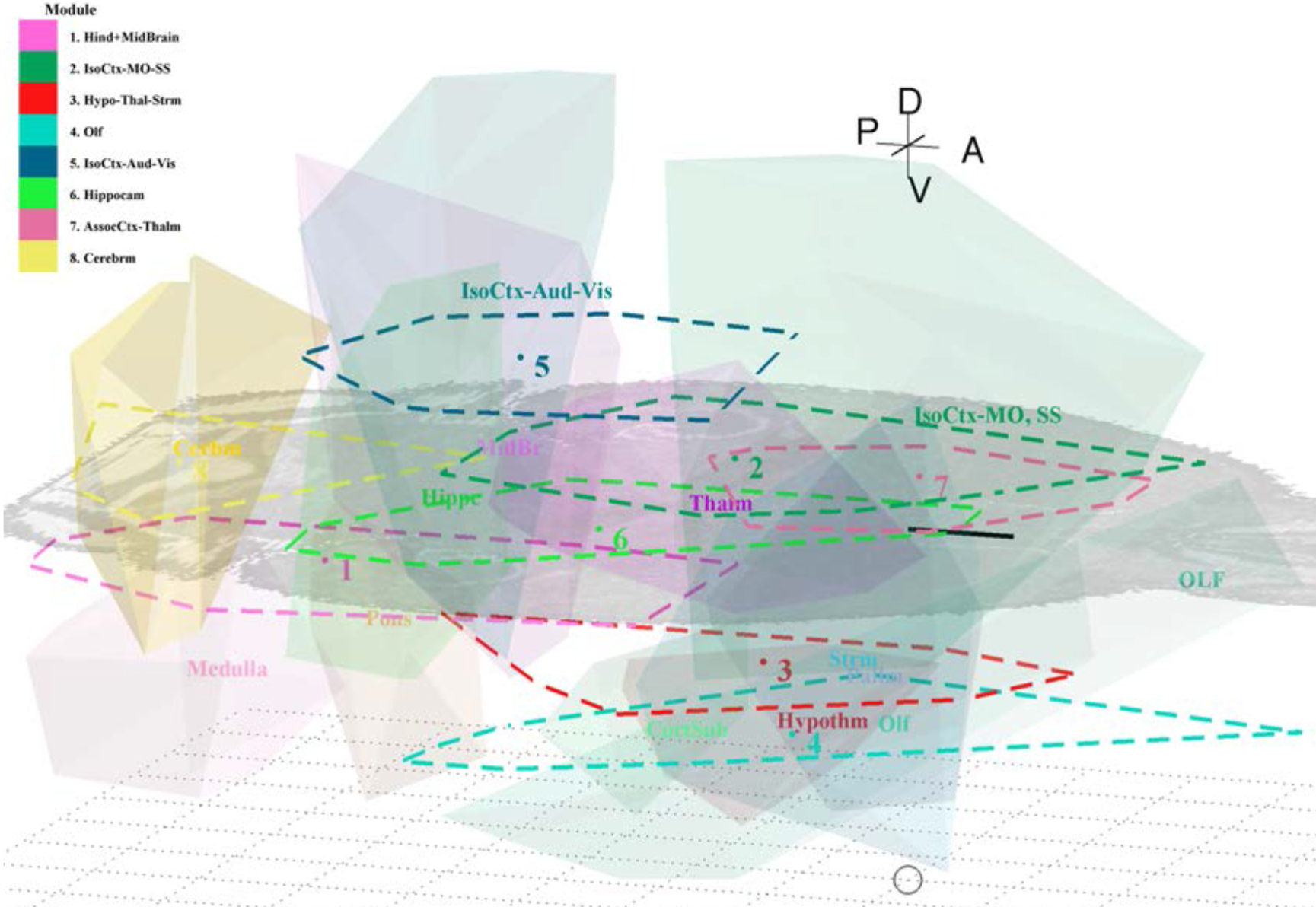
Network modules and brain regions. 3D plot of mouse brain major anatomical regions, drawn as a translucent convex hull enclosing centroids of member anatomical regions; and Infomap modules, enscribed in 2D (at height (ventral-dorsal axis) of the CM of nodes). Region colors are taken from the Allen Atlas sagittal images [63]; module colors follow that of the dominant members. Bregma is marked by the cross. A mid-level horizontal slice through the 3D Nissal stain image [14] gives perspective. A 1 mm scale bar is shown on that mid-slice. A floor grid of 1 mm squares is drawn at 6.7 mm ventral to Bregma to give further 3D perspective.

Table 1 lists the Infomap modules and their key characteristics, (cf. S2 and S3 Data for full details). Module 1 is the largest and mostly contains anatomical regions from the hind brain (medullla, pons, cerebellum) along with some (27%) from the midbrain. Module 2 is dominated by cortical (motor, somatosensory) and thalamic regions; it also contains CP, the largest and most densely connected region. Module 3 is mostly associated with the hypothalamus, along with the striatum and thalamus. Module 4 is dominated by the olfactory regions in two broad locations: anterior (MOB, AOB, AON, TT) and posterior to Bregma (PIR, COA, NLOT, PAA, TR); it includes the perirhinal region. Thus this module has a significant spatial extent in Figure 2. Module 5 contains the auditory and visual regions of the cortex, along with their thalamic pathways (LGd, MGd); it also contains the nearby, posterior cortical association areas (parietal, retrosplenial, temporal). Module 6 is dominated by the hippocampal formation, along with some elements of the palladium and striatum. Module 7 contained the anterior cortical regions (orbital, limbic, insular, cingulate) along with the claustrum and five nearby thalamic regions (AMv-d, SMT, MD, CL). Module 8 is dominated by the cerebellum. In each case a small number of nodes in other regions are well connected with the main elements of each module - so there is not a complete alignment between regions and network modules. Three regions (FS, MH, RH) appeared as “orphans” not assigned to any module – judging by their dominant linkages they appeared to be related to Module 1, so were assigned there for convenience. Modules were color coded by their dominant regional members. In subsequent figures nodes and links are colored according to their module membership, as described above.

**Table 1.** Informap modules for mouse brain ipsi-lateral links. Key members, in order of probability flow, and color coding used in the figures are listed.

| Module | Flow in Module | Key regions | Anatomical Regions | Color |
| --- | --- | --- | --- | --- |
| 1 | 0.321 | MRN, PAG, SCm | Midbrain, Pons, Medulla | pink |
| 2 | 0.229 | CP, SNr, GPe, Mop, SSp, SSs | Striatum, Thalamus, Cortex | mid green |
| 3 | 0.159 | LHA, SI, ACB | Hypothalamus, Palladium, Striatum | red |
| 4 | 0.102 | MOB, AON, PIR, ENTI | Olfactory | olive green |
| 5 | 0.068 | Vis, Aud, TEa, LGd, MGd | Cortex, Thalamus | dark green |
| 6 | 0.058 | CA1,2,3, DG | Hippocampus | light green |
| 7 | 0.038 | ACA, ILA, AIV, ORB, CLA | Cortex, Thalamus, Cortical Subplate | salmon |
| 8 | 0.022 | AN, SIM | Cerebrum | yellow |
| orphans | 0.003 | FS, RH, MH | Pons, Thalamus | n/a |

Functional roles associated with these modules emerge from analysis of sensory and association pathways, presented below. This accords with other findings of modules that broadly align with anatomical regions [68] (and cf. Figure 2). The modular decomposition found also approximately aligns with the four system model [69], that divides the brain into four functional systems: sensory, motor, cognitive, and behavioural state. Network modules 2, 4 and 5 handle sensory roles; module 2 also contains the motor roles, modules 2 and 5 contain the cortical association areas, and modules 1, 3, 6 and 8 relate to behavioural state. A key nodes (e.g. CLA, ENTl) appearing as linkage hubs between modules.

Other modular decompositions are compared in S3 Table. The Newman modularity metrics [55–56] are: Q=0.40 (Infomap), Q=0.395 (Louvain) and Q=0.30 (Newman spectral). The Louvain method produces 6 modules for the whole brain. The Infomap modules 4 (Olf), 1 (Mid- and Hind-brain) and 8 (Cerebrum) are combined together into the Louvain module 1. The Louvain results substantially overlap with those of the Infomap method, with notable differences, as listed in S3 Table). The Newman spectral partition, by definition, produces only 2 modules, essentially dividing the brain into sensory + motor + cortex and hindbrain. Thus, at a gross level, the three methods are in accordance. Other Newman methods did not produce satisfactory results, either yielding only one large module, or very unbalanced divisions. Some modules were investigated as stand alone systems, to see if further sub-divisions could be extracted. In particular the Infomap module 5 was investigated, since it combined the Auditory and Visual systems, in contrast to the Louvain method, where they are separated. This did not reveal further partitions. Infomap module 1, however, did subdivide, into groupings of Midbrain, Pons and Medulla.

A recent analysis of additional mouse connectivity, at the layer level of the cortex [12], used the Louvain method for modular decomposition. It found 6 modules with some similarities and notable differences to the present results. The 6 modules were described as: prefrontal (FRP, ACA, ILA, ORB, MOs); antereolateral (GU, VISC, AI-d, -p, -v); Somato-motor (SSp-m, -n, -bfd, -ul, -ll, -tr, MOp); Visual (Vis-p, -l, -al, -pl); medial (Vis-am, -pm, RSP); and temporal (AUD, TEa, PERI, ECT). Notable differences in the present work are that: the frontal pole, insular regions and GU are together in Infomap module 2, comprising somato-sensory and motor regions; here MOs and MOp are in this same module. The AUD, VIS and VISC regions come together in module 5, along with TEa, ECT, RSP. Five of the 8 Infomap modules (1, 3, 4, 6, 8) found herein encompass hind- and mid-brain regions along with the olfactory regions; of these module 1 includes PERI. Three modules are associated with cortical regions: module 2 (ACA, ILA, ORB, all SSp, SSs), module 5 (AUD, VIS, VISC, RSP, TEa), and module 7 (ACA, AI-v, ILA, ORB). In the present study the whole hemisphere, rather than just cortical, connectome was decomposed into modules. Thus nodes and modules are more inter-linked, including via extra-cortical regions. Whereas higher resolution, at the cortical layer level [12], may have facilitated decoupling of auditory and visual regions.

All calculations of modules based on the full adjacency matrix, independently of cut-off weight assumed, produced a two-level decomposition: no hierarchy was detected. That also was found for the mouse retina microscopic connectivity data [11]. However examination of a distance-sliced adjacency matrix did reveal an evolution of hierarchy, which was evident for closely spaced nodes whose links activate early in signalling pathways. Progressive inclusion of longer range links washes out that hierarchy, which is swamped by denser interconnections formed at later times, or equivalently longer link distances, as described below (cf. S9 Figure).

### Network Visualization

S4 Figure shows all 213 anatomical regions as network nodes plotted in 3D at the Centroid (mean coordinates) in the Allen Atlas coordinate space, along with all their Out-links. Nodes are coloured by their module membership and links by the module membership of their target. However a plot of all nodes and their In-links is not discernibly different, despite the differing Out- and In-degree patterns (cf. S2 Figure). Overall linkage patterns, as reflected by the link colors, appear to correspond with brain regions. This is consistent with Figure 2, which shows significant overlapping of network modules with anatomical regions. While there are difference in the patterns of Out- and In-links such cannot be immediately discerned in these images. This immediately highlights the basic challenge that is usually encountered in network visualization: the fabric of links is too dense to discern more than the overall pattern. In particular there are too many links crossings that occlude other features in the 3D fabric of nodes and links. As with experimental investigations of brain networks, a subset of nodes and links needs to be targeted to discern details of regions and circuits.

Figure 3 shows all nodes and the strongest 40% (5,641 links in total) of ipsi-lateral Out- and In-links with all 213 nodes, plotted in the simplified schematic coordinates. Overall there is a reciprocal pattern of links between major regions. Cortical association areas, including the ventral olfactory region (PIR), along with motor and selected midbrain regions, are significant integration hubs. From the color coding of links it appears that many of these are hubs with their relevant module (e.g. Frontal Pole, Orbital, Insular), while others are integrating between several modules (eg. Cingulate, Temporal, Parietal, Retrosplenial) or across several senses (Aud, Vis, Olf). Closer examination reveals mutual, strong (i.e. top 40%) links between somato sensory regions and both auditory and visual regions, with the temporal, parietal and cingulate regions having strong in-links from auditory and somatosensory regions. The retrosplenial region receives inputs from the gustatory, visceral and visual senses. The orbital, limbic and cingulated regions have strong In-links from the olfactory and auditory senses. The somatosensory, gustatory, visceral, auditory and visual primary senses all link directly to the motor regions. There is no direct link from the primary olfactory regions (AOB, AON, MOB) to either MOs or MOp. Possible weaker links are examined later. It is evident that a further strategy is required to unravel the complexity of links evident in Figure 3.

**Figure 3.**
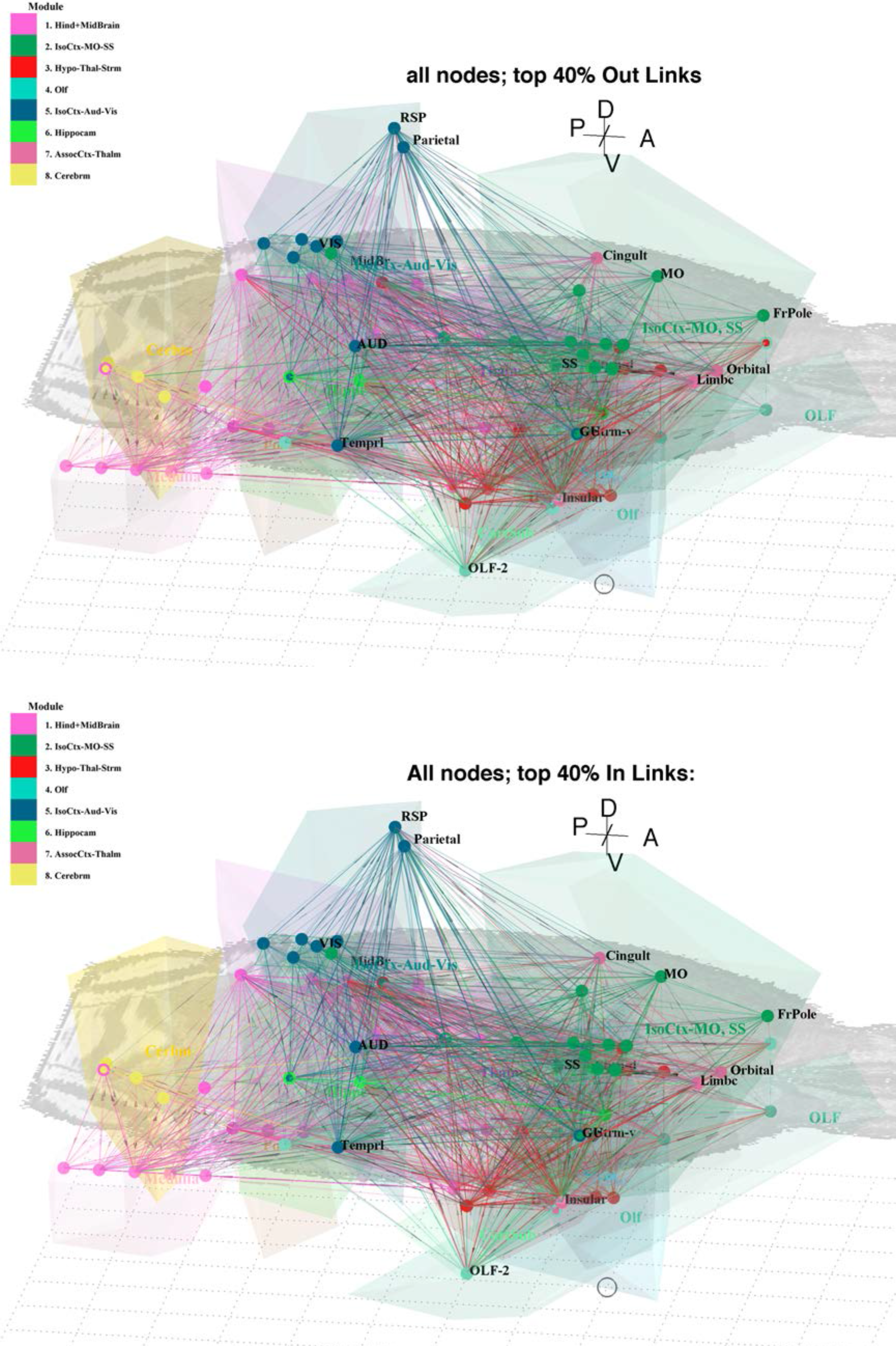
3D schematic coordinates for mouse brain. All network nodes plotted in 3D schematic coordinates (cf. S10 Figure and S3 Data), along with the top 40% of links: **a**. Out links and **b**. In links. Anatomical regions outlined as in Figure 2. Only the right hemisphere and ipsi-lateral links are shown. Other details as in Figure 2.

### Unfolding of links in sensory pathways

The network analysis present thus far considers of the linkages patterns and their spatial separation, and ignores time. That is they present the long time, steady state behaviour of the network of nodes and links. Now we examine the unfolding activation of those links through time, or over distance. The results presented above relate to nearest neighbour links, over one step, between nodes. Recalling S3a Figure it is clear that many node pairs are separated by 2 steps over the network. Likewise Figure 3b suggests that, for a given signal velocity over axons, two shorter links can be traversed in the time window for one long link to be traversed. It is also evident from Figure 1 and S1 Figure that shorter links tend, on average, to be stronger than longer links.

Figure 4 shows the top 40% of out- and in-links from/to key primary sensory regions: Visual (-p, -al, -am, -l, -pl, -pm), Auditory (-d −p −v), SSp (-bfd, -n, -m, -ul, - tr, -ll), Olfactory (MOB, AOB, AON) and Gustatory + Visceral, 20 source nodes in all. Here only ipsi-lateral links are described. These links are shown in two ranges of link, or equivalently signal time, windows: d < 5 mm (Figure 4a) and 5 < d < 10 mm (Figure 4b). The first set is equivalent to about half the brain size, being the approximate width of one hemisphere or half its length; the second almost spans the full size – in the present case no longer links of high weight were present. The 20 source nodes together have 530 high weigh direct links (top 40%): of these 426, or 80%, have link distance d < 5 mm while 104 are longer range, up to 10 mm. Two-step links begin to appear for d > 3 mm, or equivalently t > 3 msec; in the short interval (d < 5 mm) 204 such structural links are present. Of those possibly only a fraction are activated to functional roles. The second window (5-10 mm links) contains the remaining 15,367 two-step structural links from the combined 20 source nodes. The images superimpose many such links given the common structural coordinates assigned to nodes so many details are masked. Even so, some integrative features emerge: there is a clear anterior-posterior localization of processing on shorter time scales. Additional details are listed in S1 Text.

**Figure 4.**
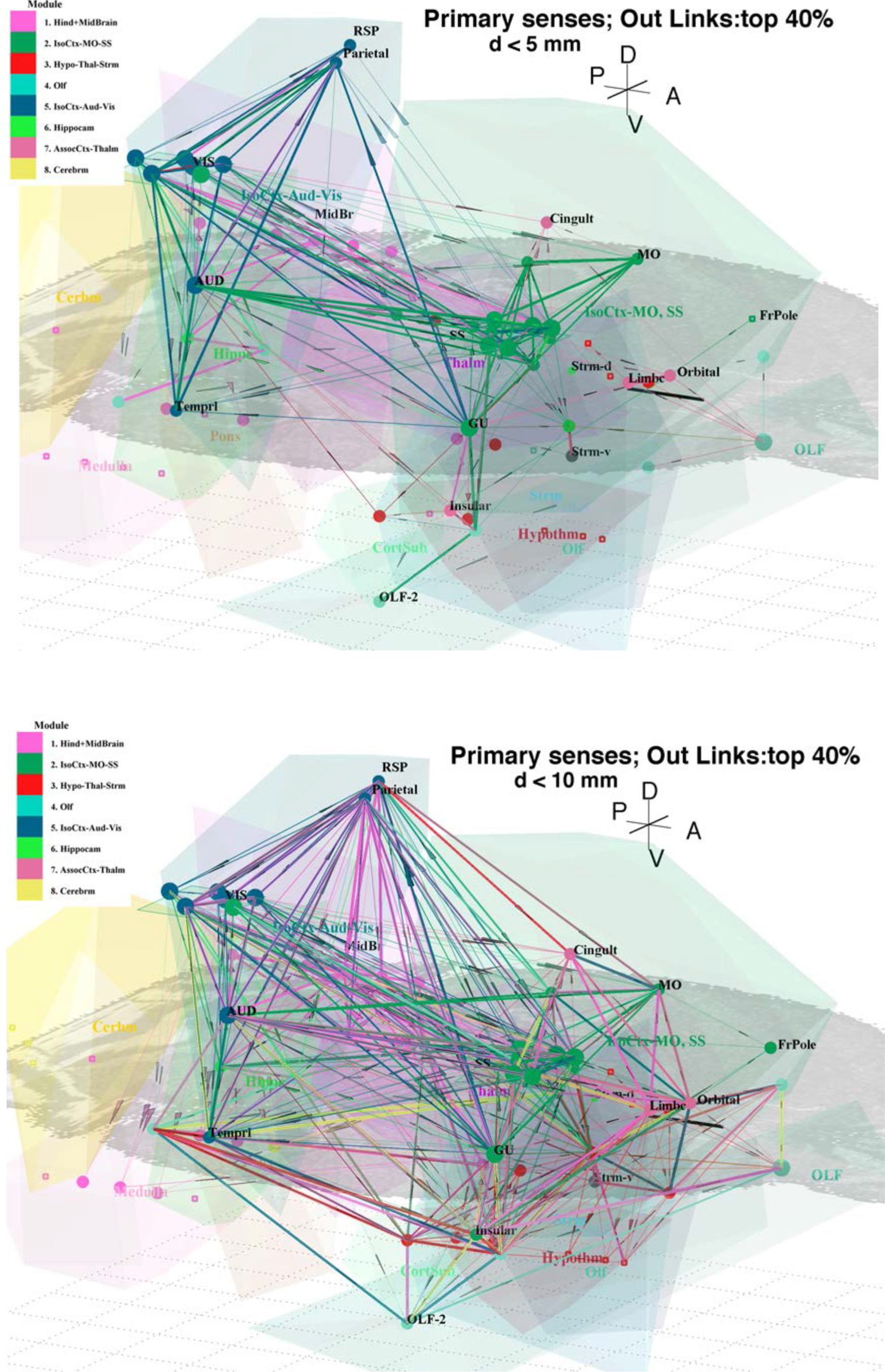
Primary sensory pathways. Major sensory pathways, shown by strong Out-links (top 40%) from 20 primary sensory areas (listed in text) plotted in schematic coordinates: **a**. d < 5mm and **b**. 5 < d < 10mm. Nodes are colored by their module membership, and links by the module membership of the target node.

Shorter links (Figure 4a) carry sensory signals to nearby association areas, e.g. Olfactory (Olf) to the nearby Limbic and Orbital regions, and then later to PIR and TT; Somatosensory (SS) regions output directly to Motor, Limbic, Orbital and Cingulate regions, along with the dorsal Striatum and the Gustatory/Visceral (GU/Visc) regions. Auditory output is direct to Temporal and Parietal regions, and also to Visual regions (Aud-d to Vis-l, -am, -pm; Aud-v to Vis-al, -am; Aud-p to Vis-p). Visual output is direct to both Parietal (from Vis-p and Vis-am) and Retrosplenial (from Vis-am) regions. Both Auditory and Visual primary regions link directly to selected Somatosensory regions (Aud-d, -v to SSp-bfd; Aud-d, -p, -v to SSp-tr; Aud-d, -p to SSp-ll; and Aud-d to SSs). Cross modal linkages are evident, e.g. between Visual and Auditory outputs, as detailed in S1 Text. At longer times (i.e. longer links in Figure 4b) anterior-posterior coordination emerges; there is more between-module interactions and cross-talk between cortical association areas. The Parietal and Retrosplenial regions link to multiple regions of the Midbrain; and many regions now link to ventral and hind brain regions. Olfactory outputs travel via PIR to the temporal cortex, to GU/Visc, to one amygdilar region (COAa) and ventral regions: Hippocampus (CA1), Cortical Subplate, Striatum and Palladium.

From the 3D images a number of regions emerge as apparent network hubs, sending or receiving multiple links between sensory modalities or between modules (indicated by link colors). Note that such regions were not immediately identified by the network “hub” measure, suggesting that a combination of techniques may be required to dissect key nodes and links and identify hubs. These emerge at link lengths, 5 < d < 6.5 mm, and most are in the Olfactory system, with each having ~20-45 In-links in the top 40% of links. S5 Figure show these links in two distance (or, equivalently, time) slices. The 5 target hubs are: ACB (nucleus acumbens, Striatum) with very strong (weight > 10k) In-links from the Insular region (AIv), Thalamus (PT, IMD), Cortical Subplate (BLA) and Midbrain (CLI) – here source nodes are listed by increasing distance from the target. Further hub-like targets are discussed in S1 Text. PERI is noteworthy in having numerous, diverse and strong outputs to seven of the 12 major anatomical regions, along with feedback to some senses (SSp-n, -m, -bfd, tr and AUDd), and a weak link (weight 100) to the frontal pole. The other “hub” nodes discussed have relatively fewer (~ 10-30) and weaker Out-links. EPd is notable: it strongly links (weight 4.7k) to ENTl – that, along with ACB, were identified as hubs via the participation coefficient (cf. S2 Table).

### Fabric of Weak Links

Figure 1 shows a strong fall off of link strength with distance: selection of the top 40% of links, while avoiding false positives, also omits some potentially important weak links. Suggestions that weakest links may have other characteristics or roles, such as stabilising complex networks [70] suggest that weaker links should not be ignored even at the risk of including false positives. Now we examine the fabric of weak links out from the primary senses, specifically the 20 nodes examined in Figure Recalling Figure 1, there are 2751 links with weight < 1 (and > 10^−6^, the weakest included herein); of there 369 links emerge from the 20 primary sensory nodes. As with S5 Figure, it is illustrative to separate links into short range (reached in shorter time) and longer range: 25% of such links have length < 5 mm, while 62% are < 6.5 mm. By contrast 80% of the strong links (Fig 4) have length < 5 mm. In the following links are listed in order of increasing distance, with shorter first. S6 Figure shows these weak links in the two distance, or equivalently time, windows. S6a Figure shows the shorter (d < 6.5 mm) weak links: note that, in contrast to the strong links, they do not target the motor regions or most of the cortical association areas. Only Cingulate and Retrosplenial regions have a few weak In-links. All weak links, up to length < 10 mm, are displayed in S6b Figure. Now there are multiple long-ranged feedback links to the Cingulate and Retrosplenial regions, and a number of possible two-step links to Motor regions: 111 into MOp: eg. MOs – prelimbic – MOp, MOs – Cingulate (ACAv) – MOp, SSp-m – STN – MOp, etc; and 128 into MOs: eg. AON – TRS – MOs, SSp-n – TRN – MOs, MOB – TRN – MOs, etc. It is unknown which, if any, of these very weak structural links are functional. There are no one-step weak links into MOp or MOs. Otherwise most Out-links from the primary sensory regions are to the behavioural state system (in Cajal’s classification [69]), the hind brain, thalamus, hypothalamus, and hippocampus, along with the midbrain and ventral olfactory regions. There also are 15 very long range (10 < d < 12 mm), weak links (not shown), the majority from MOB, the most anterior anatomical region.

### Cortical Association Areas and Sensory Integration

The combination of network analysis and visualization also permits examination of concurrent In- and Out-links to the cortical association areas. The separation of the Isocortex into anterior (Somatosensory-Motor) and posterior (Auditory-Visual) modules (Figure 2) suggests examination of the two regions separately. The four anterior association areas: Insular, Limbic, Orbital and Cingulate are shown in Figure At first sight the In- and Out-links appear to be symmetric. However closer examination reveals detailed differences, as discussed in S1 Text.

There also are 399 strong (i.e. top 40% by weight) Out-links (Figure 5b) from these four regions. Again these are shorter range links, with 70-80% being less than 5mm, and the most probable link length is 2.5-3.5 mm. There are numerous strong local links between the four anterior association areas and all connect to the Motor regions. Further details are presented in S1 Text. Inspection of Figure 5 suggests that the In- and Out-links with anterior regions are reciprocal. However the details (cf. S1 Text) reveal some differences. In particular, there are many outputs from the four regions to the Midbrain, but few inputs. The Cingulate regions have outputs to the Temporal regions, but no strong inputs. Note that the Limbic region outputs to the Motor regions but has no strong inputs from there. The Frontal Pole has strong inputs from Limbic, Insular and Cingulate Regions, but only outputs to Orbital and Insular Regions.

**Figure 5.**
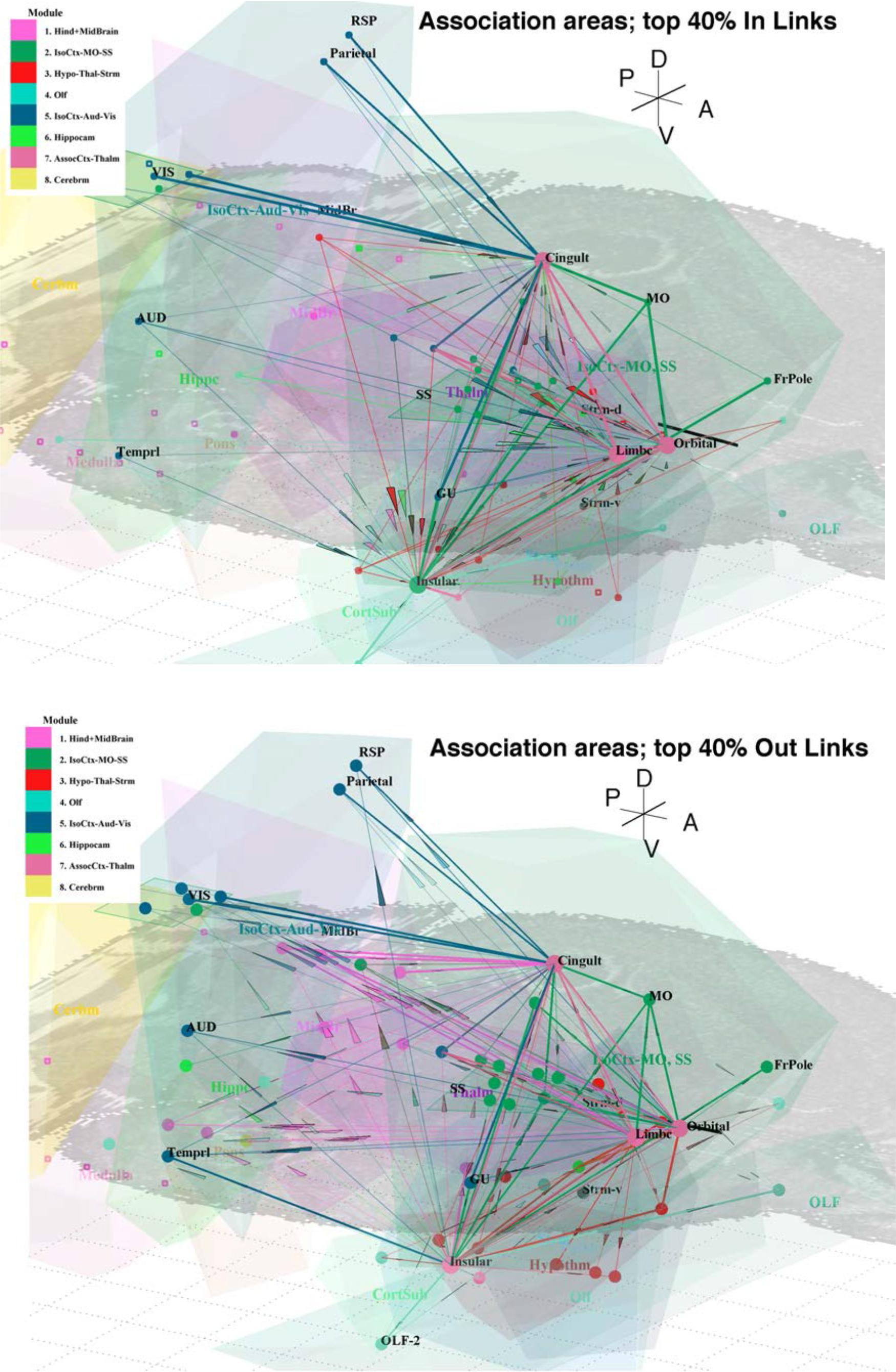
Anterior cortical association areas. **a**. Strong In-links to anterior cortical association areas (Limbic, Orbital, Insular and Cingulate). Only direct (first nearest neighbour) links are shown for clarity. Stronger or multiple links are highlighter by thicker lines. **b**. Strong Out-links.

Links with the three posterior association areas, Temporal, Parietal and Retrosplenial, are shown in Figure 6. The 216 strong (i.e. top 40% by weight) In-links are shown in Figure 6a. The figure shows that now there are longer range inputs, with temporal and retrosplenial regions having most probable links lengths of 5.5 mm, compared to 3.5 mm for the parietal region. The nearby auditory-visual and visceral regions provide immediate strong inputs to the temporal areas. The Temporal regions receive multiple inputs from the Hippocampus, very strong Thalmic input, and weaker long range inputs from the Frontal Pole. Further details are listed in S1 Text.

**Figure 6.**
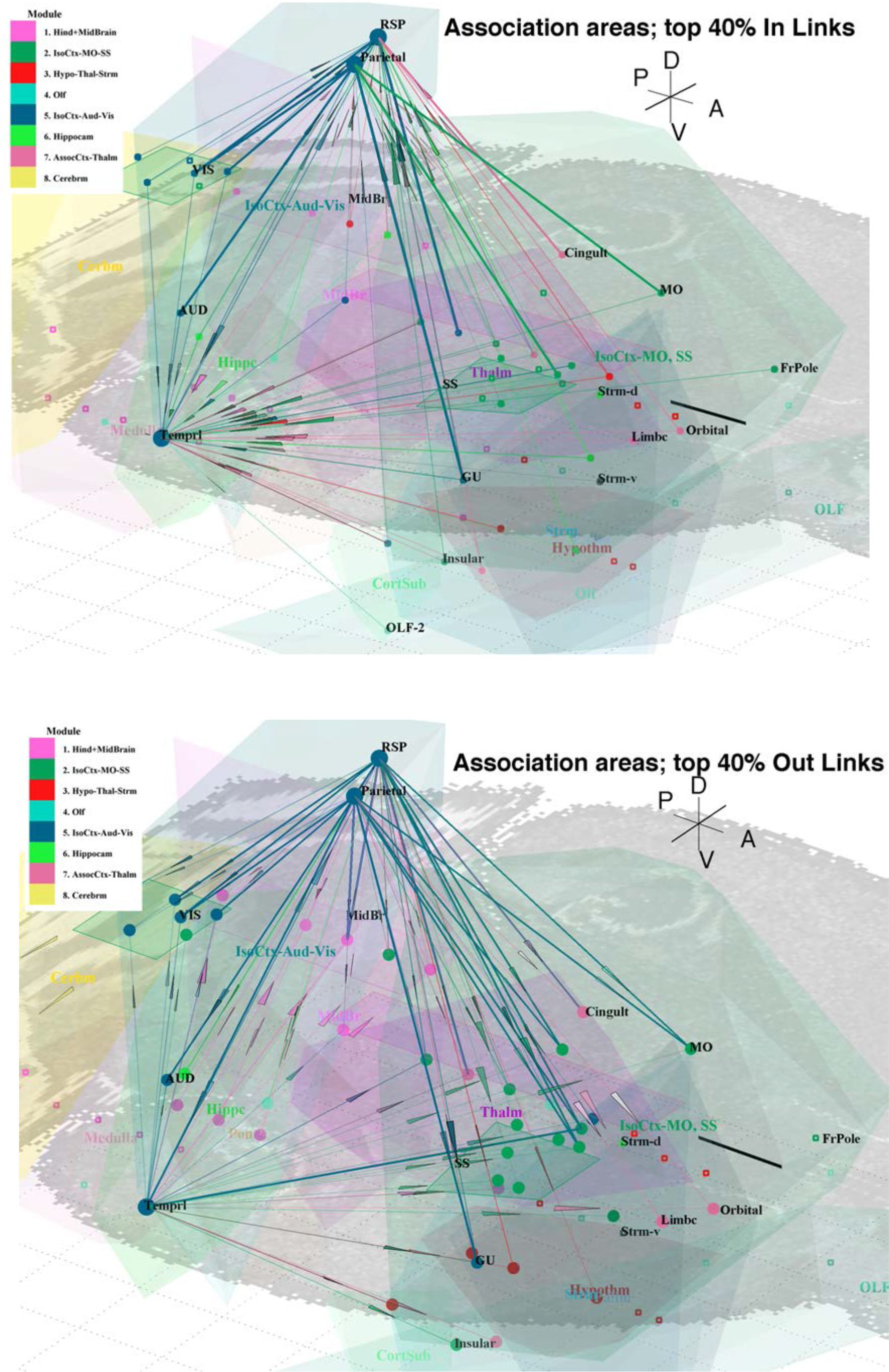
Posterior cortical association areas. **a**. Strong In-links to posterior cortical association areas (Temporal, Parietal and Retrosplenial). Only direct (first nearest neighbour) links are shown for clarity. Stronger or multiple links are highlighter by thicker lines. **b**. Strong Out-links.

There also are 204 strong (i.e. top 40%) Out-links (Figure 6b) from these three regions. Most are shorter range links, with 75-85% being less than 5mm, and the most probable links length being 3.0-3.5 mm. There is immediate feedback to the local sensory regions, numerous links to Midbrain regions, weak links to Thalmic regions, and direct links to Motor regions. Additional details are listed in S1 Text. Inspection of Figure 6 suggests that the In- and Out-links to posterior regions are reciprocal. Again the details listed above show the differences. In particular, the Frontal Pole has medium strength links to the Temporal region, but only relatively weak return links (eg. FRP to TEa, weight 37 – not shown, since not in top 40%). Similarly there are asymmetries in inputs and outputs with the Motor regions. Overall it is evident that the temporal area is a significant integration hub for sensory inputs, while the retrosplenial and parietal areas are distribution hubs to the motor areas.

The Frontal Pole is examined separately since it has more long range connections, with the most probable links length being 6.5 mm. S7a Figure shows the 53 strong (top 40%) Out-links and 17 In-links with the Frontal Pole (FRP). The closest strong inputs to FRP are from the Orbital and Insular regions, along with MOs. Strong outputs from FRP are to the Insular regions and Motor region, MOs. Other strong links and further details are discussed in S1 Text. S7b Figure shows the weaker links (bottom 60% or weight < 78) to FRP. Such have been ignored so far, for clarity and the higher chance of either False Positives or False Negative indications of functional links. Most weak outputs are to the Mid- and Hind-brain regions: Striatum, Hypothalamus, Hippocampus, Cortical Subplate, Midbrain along with the Thalamus.

### Motor Regions

S8 Figure show the strong (top 40%) links into the Motor regions (MOs and MOp). Many of the cortical association areas have such links into the motor regions: the Frontal Pole (FRP) has the strongest links: of weight 27k into MOs and 22k into MOp. Related close, strong inputs are from the Cingulate, Orbital, Insular, Retrosplenial and Parietal regions; with longer range input from a Temporal region; and direct inputs are from multiple primary sensory regions. Details are listed in S1 Text. There are also long range, weaker inputs from Pons and Medulla. Note that a number of weaker two-step structural links are shown, via the Restrosplenial, Cingulate and Limbic regions. Should they be functional they would be co-incident to MO with the other inputs. Nearby outputs from both MOp and MOs are feedback to all SSp areas and to Vis-al and, and then numerous medium strength links to thalmic areas, as detailed in S1 Text.

### Evolution of Module structure as Links are added

All of the results presented reflect the steady state modular structure of the mouse brain network, that is formed by all links present. Overall, shorter links are stronger and longer links are weaker (Figure 1). This is reflected in the sensory (Figure 4 and S5 – S7 Figures) and association (Figures 5, 6) circuits having a different character when short or long links are included. Given a constant axonal signalling velocity, this is equivalent to a time evolution of links, eg. following a sensory input event. Similar effects will determine the modular decomposition of the network when viewed through the lens of close vs. distant links. S9 Figure shown the modular decomposition of the mouse brain network calculated using the Infomap method at three stages of link evolution. Nodes are plotted now only in 2D for clarity. The first stage, shown at bottom, is formed by only short range links, of length d < 2 mm (1844 links). At this early stage there are three modules: the anterior module of 23 nodes joins the long term modules 2 (Somato-Motor) and 4 (Olf). The middle module of 74 nodes corresponds to module 5 (Aud-Vis), while the posterior module of 116 nodes comprises the ultimate modules 1, 3, 6, 8 (medulla, pons, mid-brain, hypothalamus, hippocampus, cerebrum). At this early stage hierarchy is present: the anterior module separates Motor + Cingulate, Somatosensory, and Olfactory regions, amongst others, into 7 sub-modules. Similarly the middle module separates Auditory + temporal, Visual + Parietal, and Retrosplenial regions; while the posterior module separates into 7 sub-modules.

The intermediate stage of module formation, including the 12,145 links with d < 6 mm (about half the size of the brain), is shown as the mid-level in S9 Figure. Now the early stage modules are splitting into what is almost the final modular structure. This is driven by the inclusion of many more links between nodes. The vertical lines in the figure show the splitting pathways of the initial modules. Nodes are now colored by their ultimate module membership. The final stage, including all 14,103 links (ignoring weight < 1), is shown at the top. In summary, hierarchal modularity is only evident when only short range links are included. Many of those sub-modules become separate top-level modules when longer range links are included.

### Signal flow along cortical gradients

Correlations amongst multi-modal cortical gradients [71] have been suggested as a brain organising principle, consistent with the concept of sensorimotor to transmodal spatial gradients [72]. Figure 7 shows the random walk forward transition probability, Pr(i:j) along successive steps of the SSp-bfd signalling pathway in the cortex, along with the MRI relaxation time ratio, T1w:T2w gradient along the same pathway. In each case there is a clear gradient, reminiscent of the spatial reference map previously found [71].

**Figure 7.**
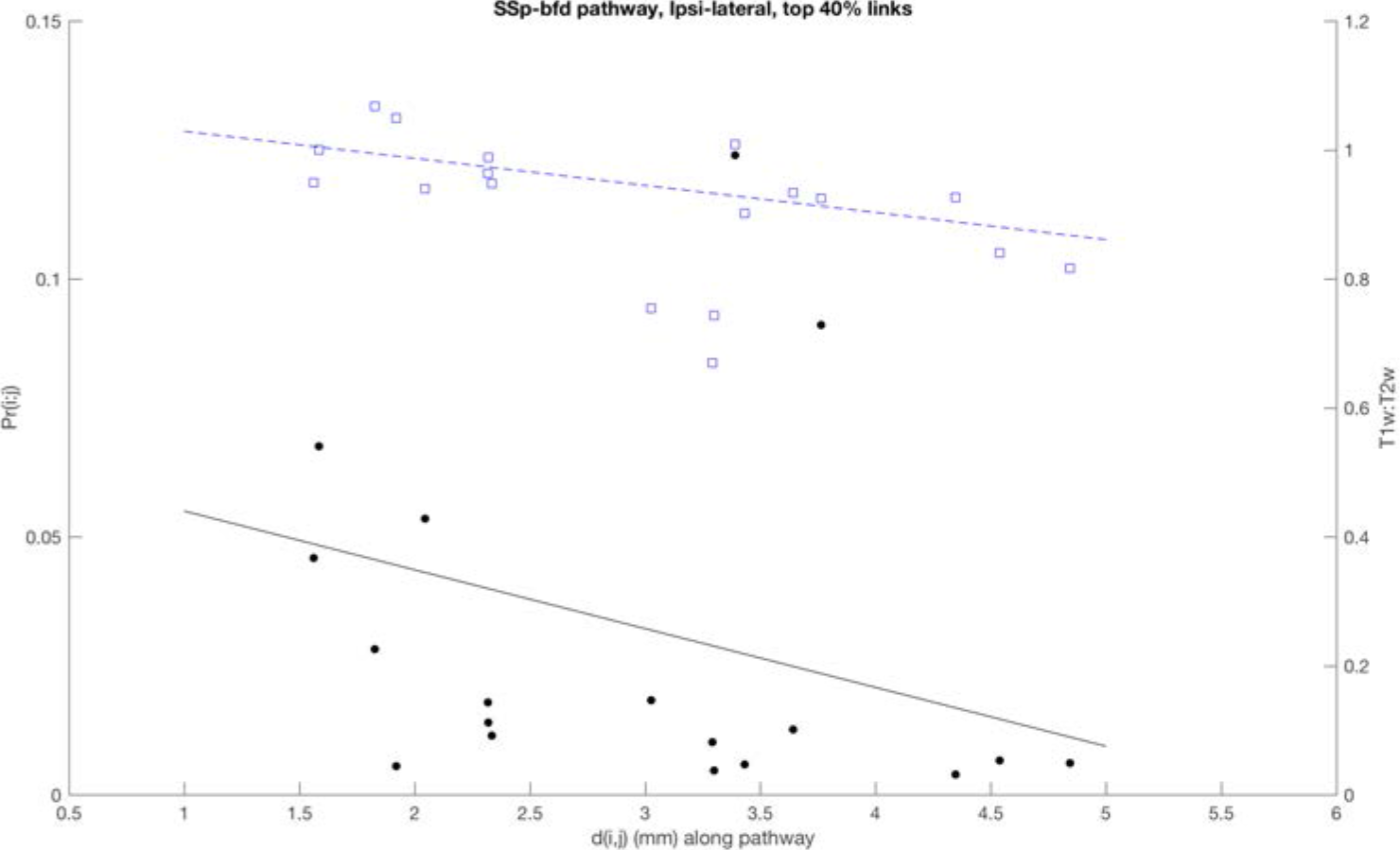
Cortical gradients along signal pathways. The node-to-node transition probability, Pr(i:j) plotted along the SSp-bfd sensory pathway in the cortex vs. the path distance (left axis and black filled circles; first neighbors only shown); the MRI relaxation time ratio, T1w:T2w in cortex vs. the path distance (right axis and blue squares). The best linear fit to each set of points is shown.

## Discussion

The Allen structural mouse connectome provides a rich data set to augment classical anatomical studies and from which to launch a more detailed investigation of brain networks and circuits. This can complement the many experimental probes that continue to provide details of specific pathways and circuits. The present analysis shows that some biologically relevant features can emerge from brain wide network analysis. Use of the full region-to-region connectivity data was justified (cf. Methods and S1 text) for most source regions. The connectivity data show a clear weight-distance fall off. This indicates that thresholds, to minimise false positives, may selectively penalise longer links, some of which may have functional significance. On average shorter links are stronger, while longer links are weaker (Figure 1). This partition shows up in the links from primary sensory regions to local cortical association areas, versus the longer range links between association areas and to the hind-brain state-keeping regions.

Simple network measures (Degree, nBC, eBC) alone do provide a sufficient guide to understanding overall network structure or important circuits. Such measures are based usually on graph shortest paths and so may be an incomplete guide to biologically important nodes or links. This is likely a symptom of the simplicity of the topological graph as a network model. Adding physically relevant features, such as spatial embedding, time evolution of links carrying fixed velocity signals, and including secondary paths for signal spread on the network, does provide a more informative model of the mouse brain network. However the participation coefficient highlights some nodes (eg. CLA, ENTl, and possibly: CLI, PP, GPi, MGd, MOs) that turn out to mediate key linkages between network modules. A judicious selection of key nodes and links, informed by known biological functions, with a focus on sensory pathways and cortical association areas can reveal features of the mouse brain structural network consistent with evidence for biological functions. Tracing the primary sensory output paths over the network reveals selective roles of the various cortical association areas of the mouse brain shows local processing, and then progression on to the motor system.

Modular decomposition based on random walk (i.e. probability) flows over the network produces plausible division of network structures devoted to primary sensing, motor activity, information integration, and state maintenance (Figure 2). This is consistent with three views of the brain: as divided into the classical anatomical regions, as four inter-linked functional systems [69], and now as comprising 8 network modules. Three of these are aligned with primary sensory pathways: Olf (module #4), SSp+MO (#2), Aud+Vis (#5); two with the cortical association areas, separated into anterior (#7) and posterior (#5) locations; and the combined mid- and hind-brain regions (#1, #3, #6, #8). No hierarchy is detected at the resolution of linkage data currently available.

### Sensory Pathways

The combined primary sensory pathways (Figures 4–6 and S5, S6 Figures) illustrate the direct, local and strong links to immediate targets, particularly in the motor and cortical association areas. The methods used herein show that the mouse isocortex can be seen in two parts, each with localised functions. The anterior senses (olfaction and somatosensory) align with the more anterior association areas: Insular, Limbic, Orbital and Cingulate. The more posterior senses (auditory, visual) interact more locally with the Temporal, Retrosplenial and Parietal association areas. Both anterior and posterior association areas have direct links to the motor regions. Secondary, in the sense of weaker and longer-range, links are between these two major regions and also including the Frontal Pole. Many cross modal structural links are present between the primary sensory regions.

The time progression of signals along sensory pathways reveals the roles of shorter vs. longer links (Figure 4). Close links are to the local cortical association areas and cross modal, while longer links provide anterior-posterior coordination and inputs to other brain regions. Figure 4b shows a number of nodes that appear as linkage hubs, some of which may have functional significance. The succession of network links from the primary sensory regions is consistent with observations of serial information flow from barrel field sensory to motor areas in a tactile decision task in mice [73]. The fabric of weaker structural links (weight < 1) is longer range and communicate with regions across the whole brain. On average the shortest path between any two nodes requires more steps (4.4 vs. 2) than over the full network with its large number of stronger, and generally shorter, links. The direct weak links are more to the hind-brain and the behavioural state system components (S6 Figure). Cross-modal sensory links are common, as described above. These strong, close cross links between auditory and visuals pathways are consistent with observations of cross-modal attention and task choices in modulated audio-visual stimuli [74].

### Cortical association areas

There is a clear anterior-posterior network locatisation of cortical association areas, which are located close to relevant primary sensory regions. They tend to have direct strong links from nearby primary senses (Figures 5, 6) and outputs to other association areas and the motor regions. The frontal pole has very strong reciprocal links locally to Orbital, Insular and Motor regions; and is distinguished by having additional longer range connection (S7a Figure) to other brain regions. FRP also has numerous weaker and wide ranging links (S7b Figure).

### Signal flow along cortical gradients

Correlations amongst multi-modal cortical gradients [71] have been suggested as a brain organising principle [72] that may provide an intrinsic coordinate system for human cortex [75]. The network description of signal pathways (Figures 4, S5, S6) is consistent with this view. Preliminary results here (Figure 7) show evident gradients along the mouse barrel field (SSp-bfd) pathway and justify further studies. Along that pathway short range near neighbor links have a higher forward transition probability compared to longer range links. There appears to be a signal flow gradient consistent with the T1w:T2w gradient has been previously suggested as a common spatial reference map [71].

### Conclusion

Many layers of network features could be significant for brain function, as discussed in the Introduction. The question is which are necessary to reproduce, and even to predict, biologically relevant features. Random walks on the full structural linkage network produce modules consistent with anatomical and brain system functions [58, 65, 69]. Biologically relevant hubs (eg CLA, ENTl) do appear as key linkage nodes identified by the participation coefficient and confirmed by the visualizations (Figures 3–6 and S5 Figure); other suggested hubs take a combination of indicators to identify as such. This highlights possible limitations of the network approach [76] at this time, and the need to reformulate network models. Ultimately higher resolution will be also be required in both the connectivity data and the level of detail included in the network models.

## Supporting information

MouseBrain_S1text

## Supporting Information

**S1 Data**. Weighted, directed adjacency matrix for the 213-node mesoscale ipsi-lateral mouse brain connectome (160 KB CSV text file).

**S2 Data.** Details of all 213 nodes along with network measures (23 KB CSV text file).

**S3 Data.** Schematic coordinates of the 213 nodes (62 KB CSV text file).

**S1 Figure.**
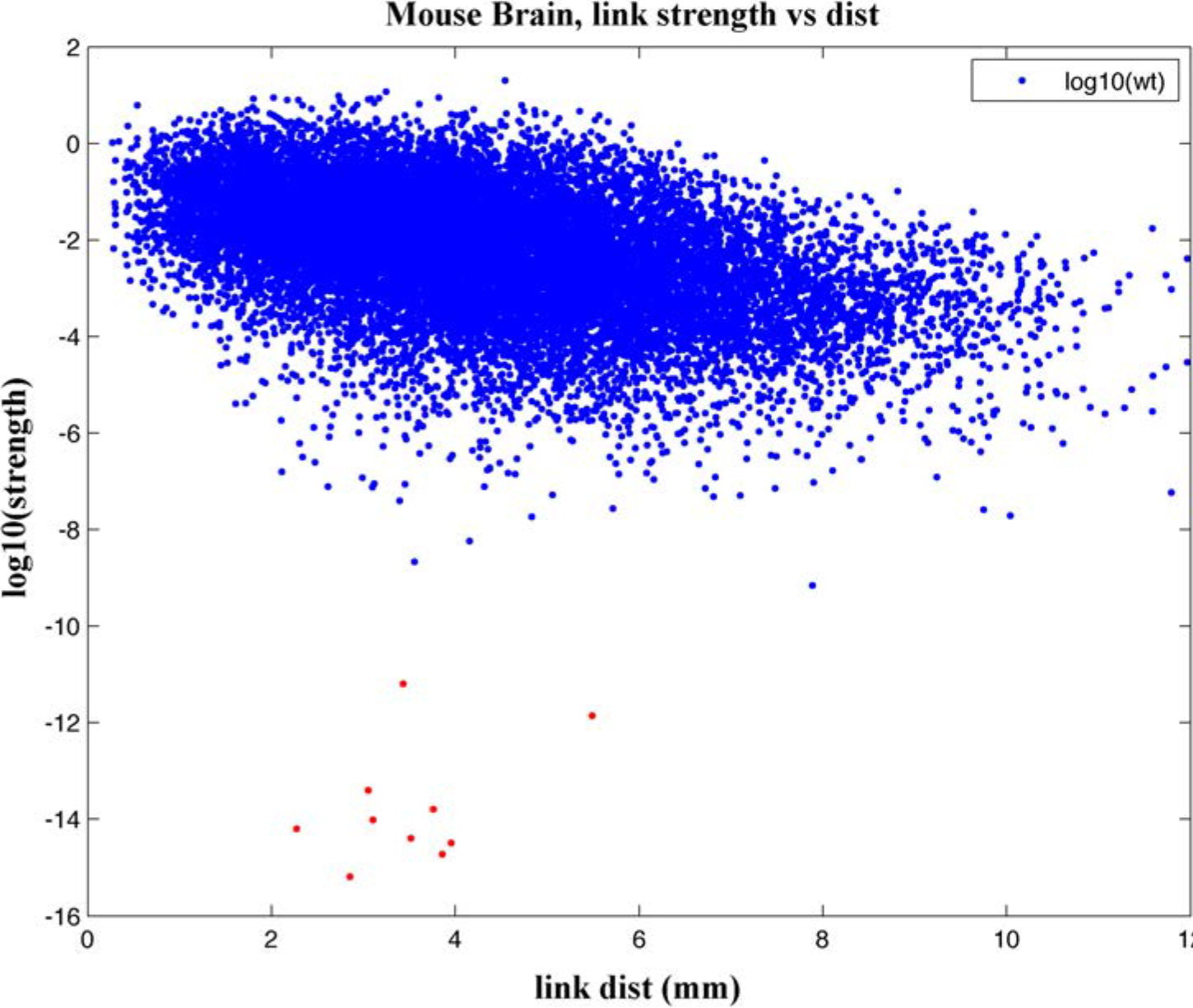
All experimentally detected link strengths plotted as log_10_(strength) vs. distance. The strength used is the quantitative projection strength reported in the original Supplm. Table 3 of [9]. The plot highlights the 10 lowest outliers (red).

**S2 Figure.**
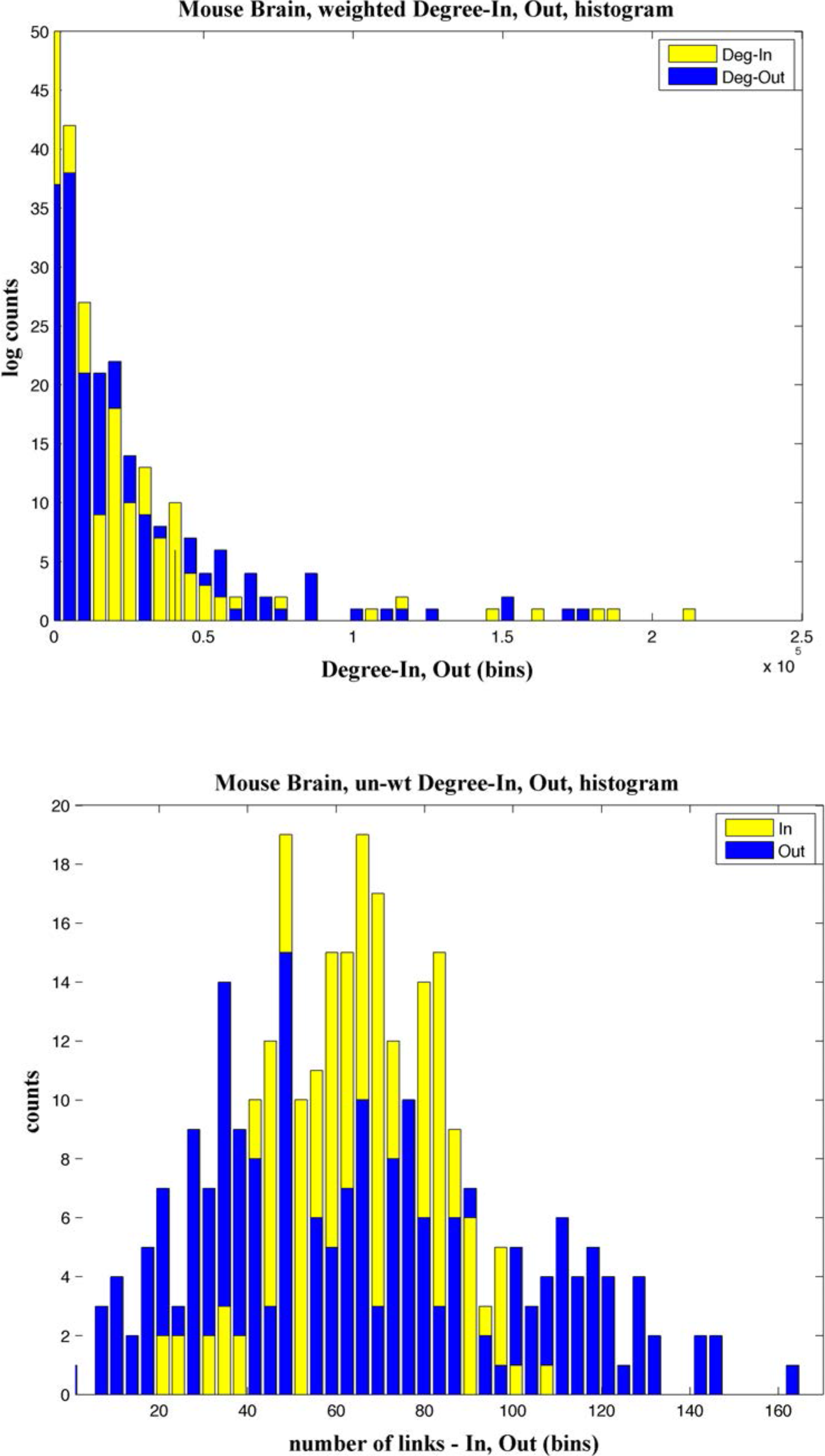
**a**. Distribution of weighted Degree-In and –Out for all 213 nodes. **b**. Distribution of un-weighted Degree-In and –Out, i.e. the number of links in and out of each node.

**S3 Figure.**
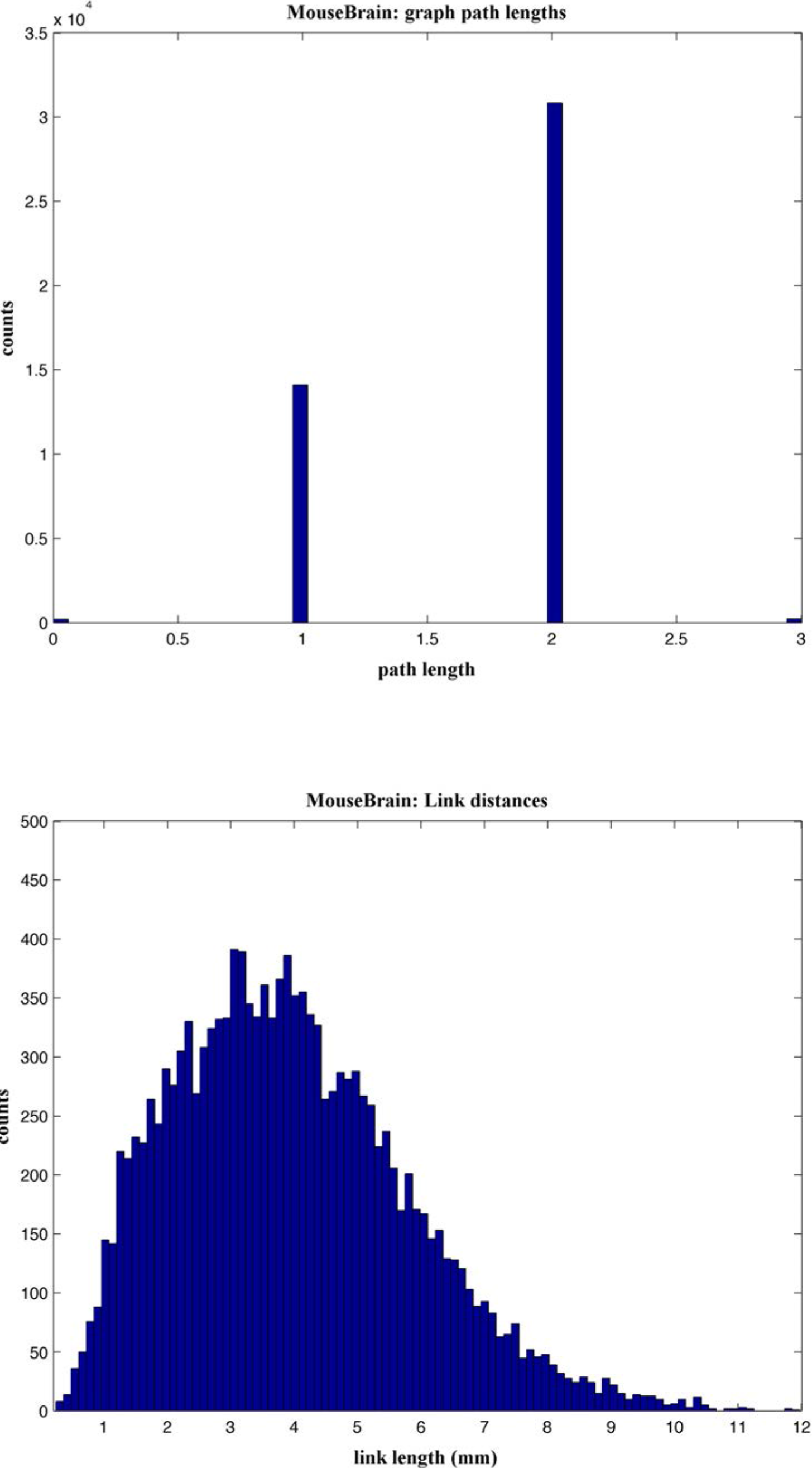
**a**. Distribution of graph path lengths (i.e. number of link between any two nodes) and **b**. Distribution of link distances in geometric space.

**S4 Figure.**
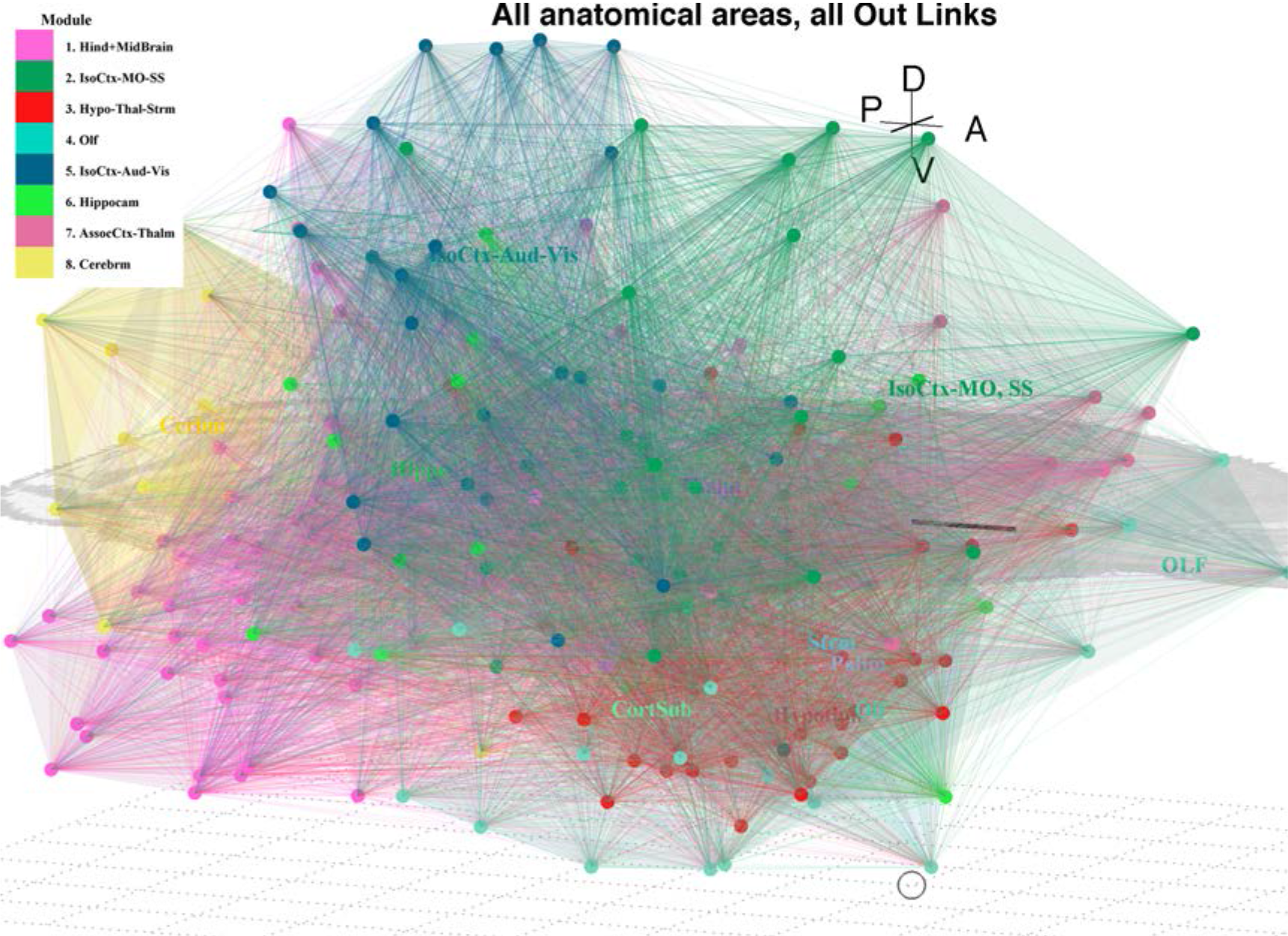
All nodes (anatomical areas) plotted in Allen coordinates and their Out-links. Nodes are colored by their module membership (cf. Table 1 and S2 Data), and links by the module membership of the target nodes.

**S5 Figure.**
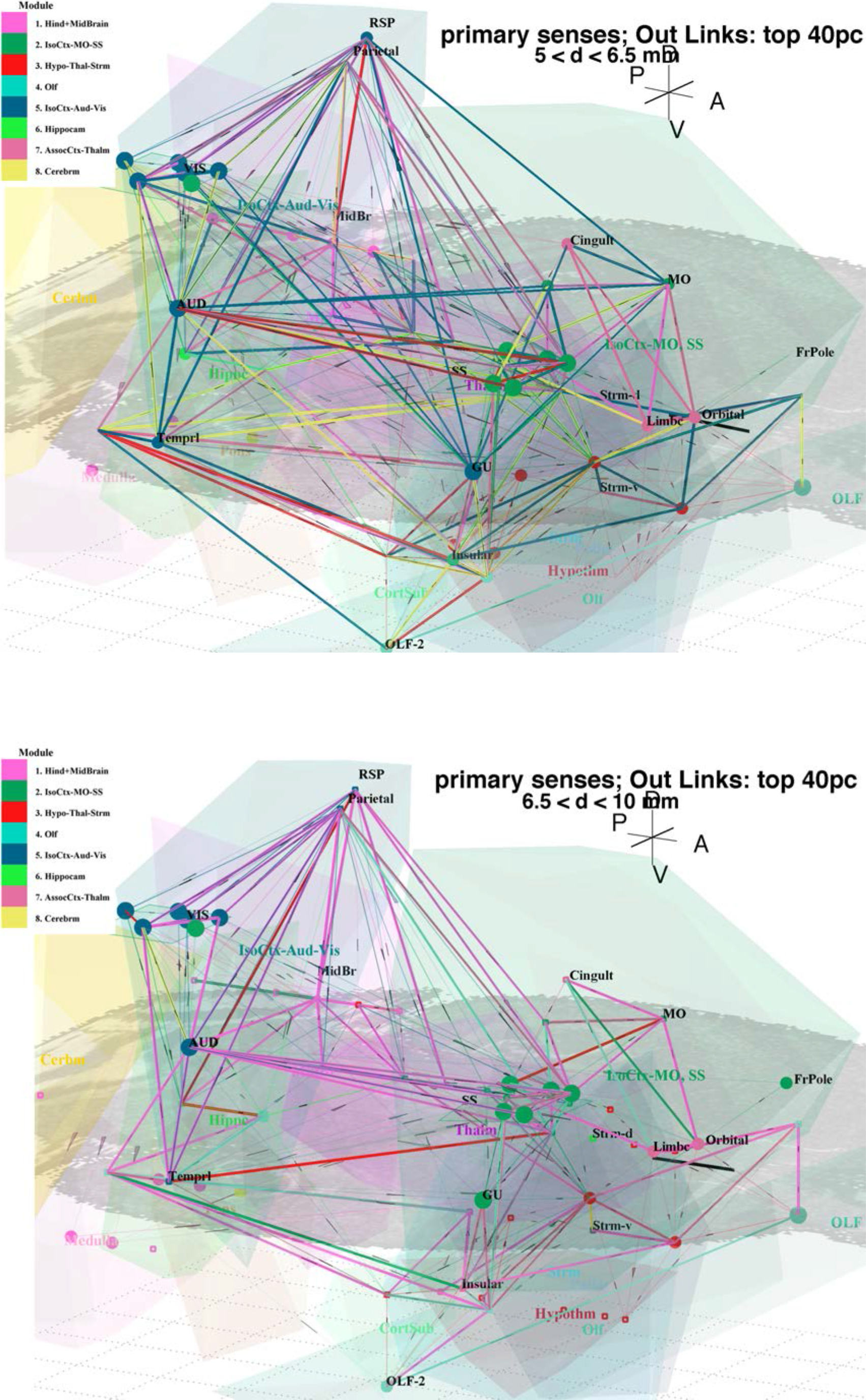
**a**. Strong out links (top 40%) from primary senses (listed in text; cf. Figure 4) for medium length links (5 < d < 6.5 mm); and **b**. for longer (6.5 < d < 10 mm) links. Shorter links (d < 5 mm; cf. Figure 4a) are omitted for clarity. Visually evident hubs in mostly ventral regions include, from right to left: DP, ACB, CP, EPd and (posterior) PERI, as described in the main text.

**S6 Figure.**
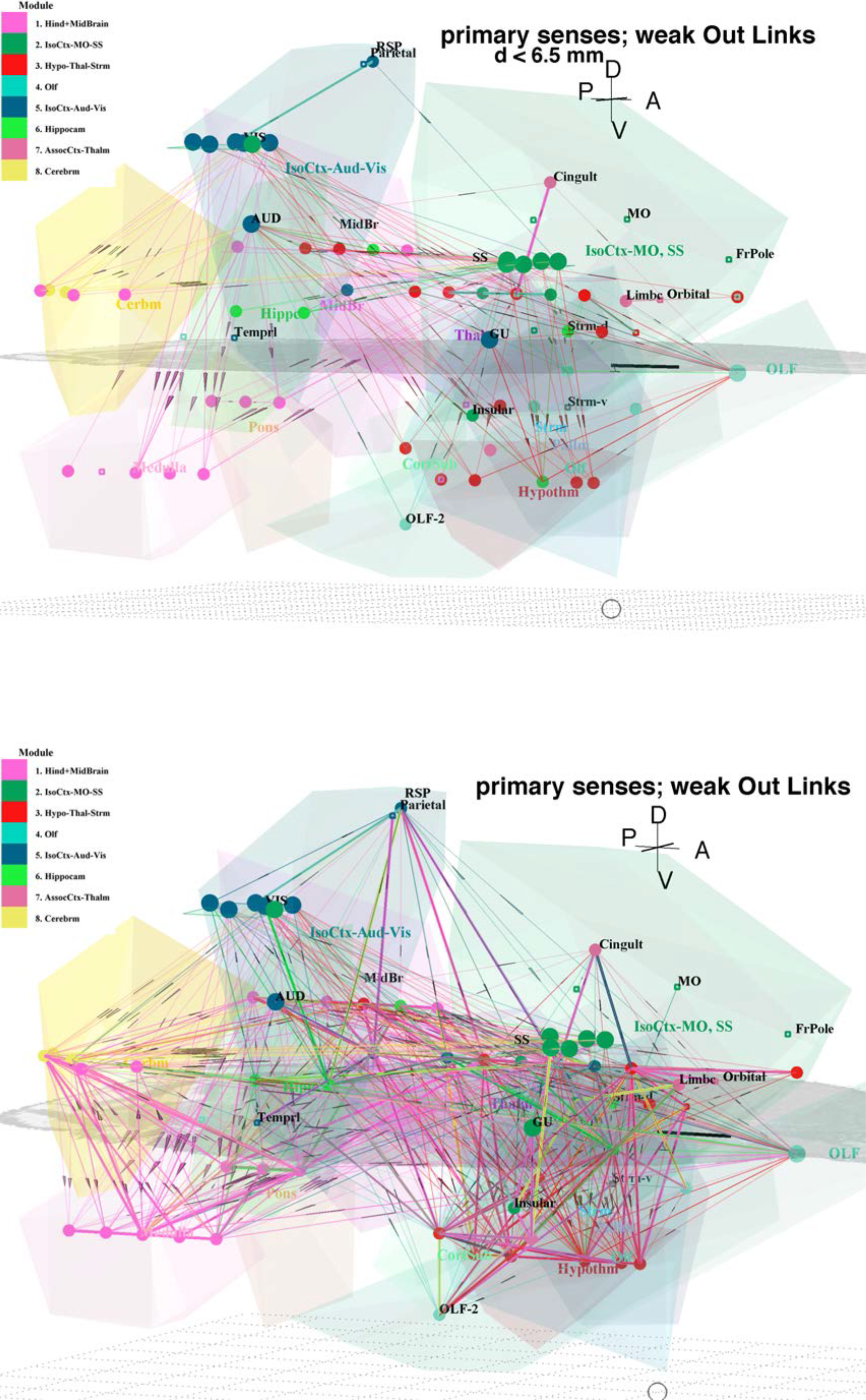
Weak Out-links (weight < 1) from primary sensory areas (cf. Figure 4) plotted in schematic coordinates, for link lengths: **a**. d < 6.5mm; and **b**. d < 10mm. Nodes are colored by their module membership, and links by the module membership of the target node. Heavier lines are caused by overwriting multiple links along the same paths.

**S7 Figure.**
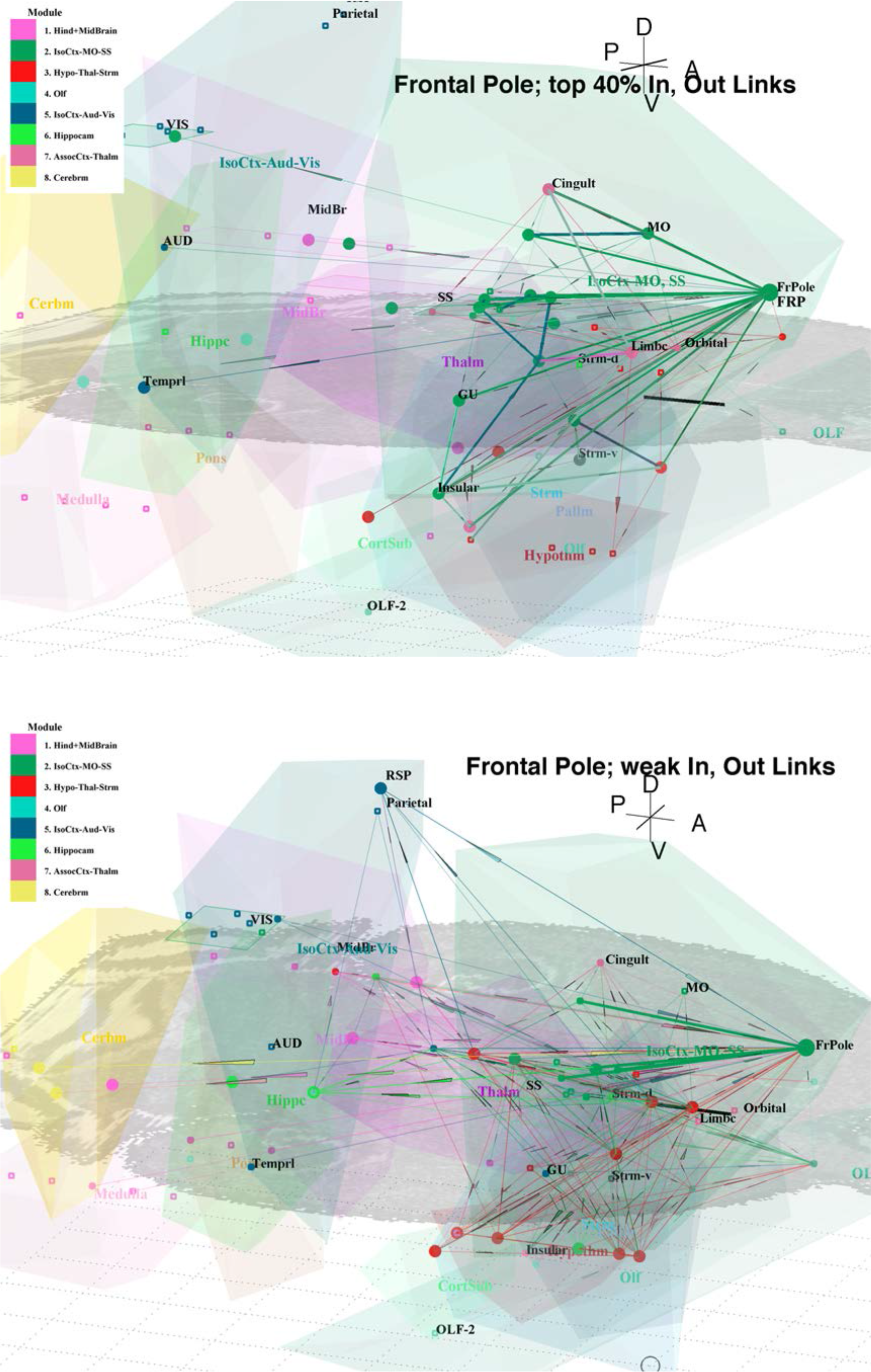
**a**. Strong (top 40%) In- and Out-links with Frontal Pole. Stronger links (weight >1 and > 5k) are highlighted by thicker lines. **b**. Remaining weak In- and Out-links with Frontal Pole. Nodes are colored by their module membership, and links by the module membership of the target node. Note a number of 2-step links are possible.

**S8 Figure.**
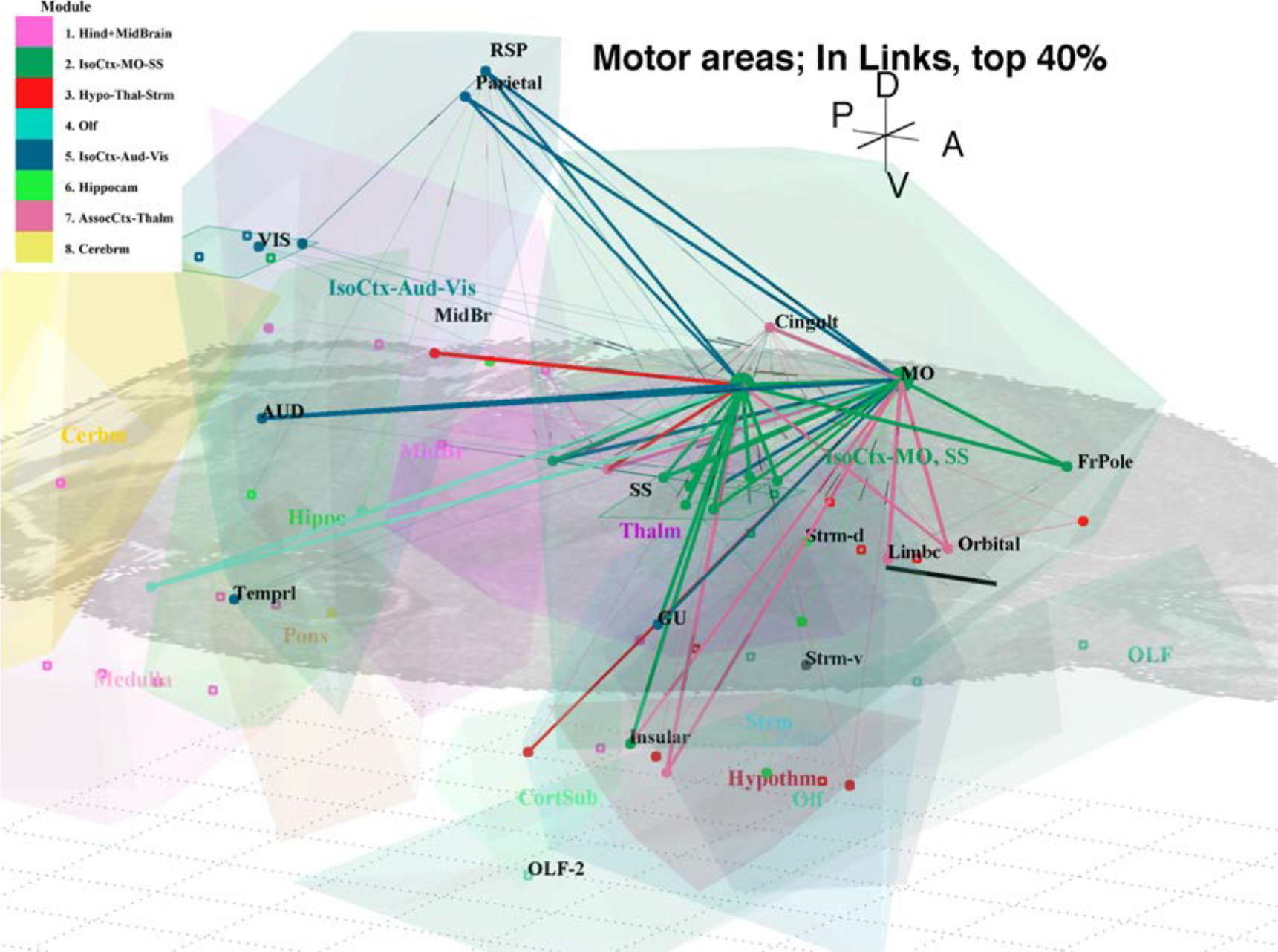
Strong (top 40%) In-links to Motor areas (MOp, on the left, and MOs, on the right – both highlighted as larger circles), plotted in schematic coordinates. Stronger links (weight >1 and > 5k) are highlighted by thicker lines. Links are colored according to their source module.

**S9 Figure.**
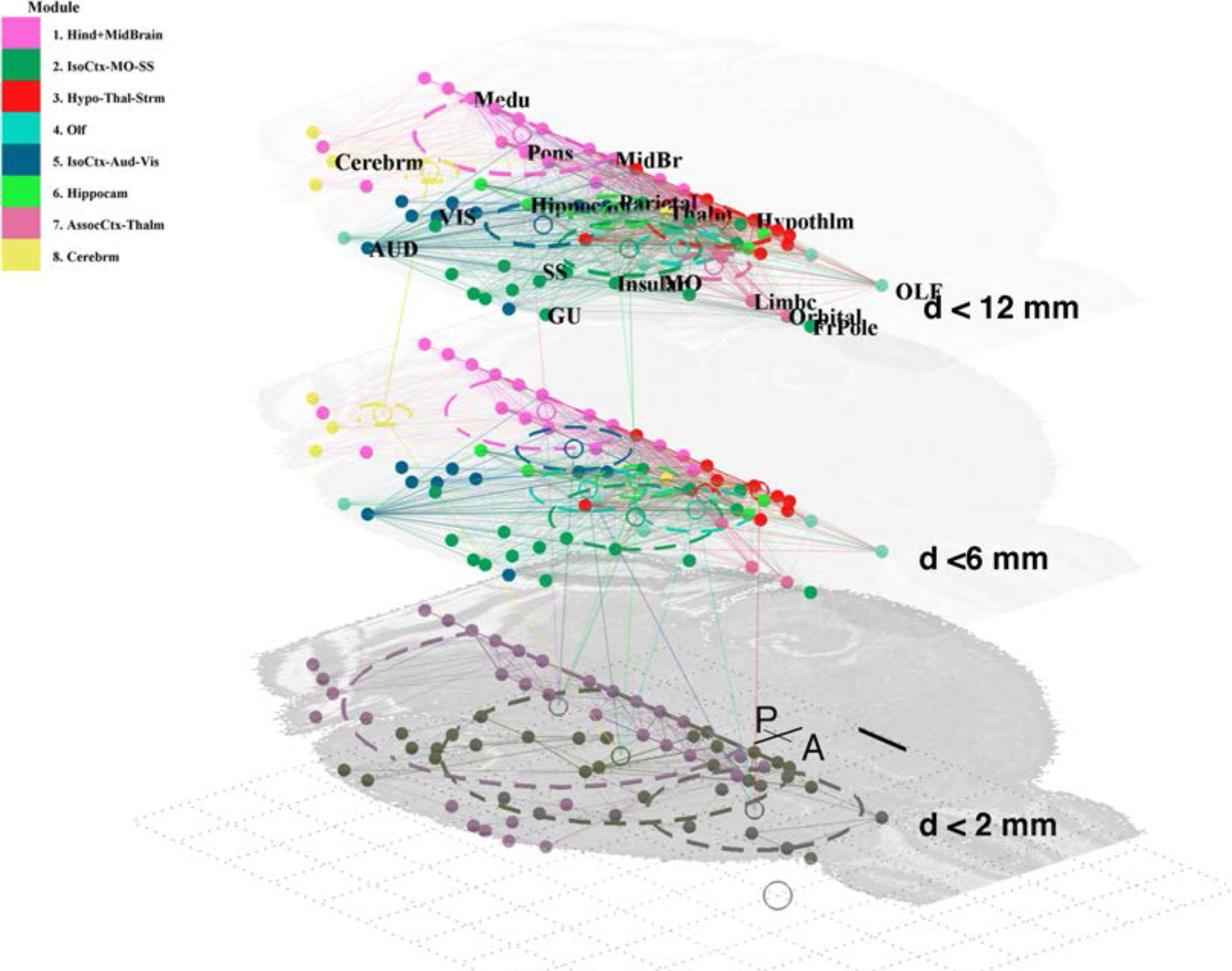
Evolution of modular structure in the mouse connectome. Nodes are plotted in schematic coordinates condensed into a 2D plane. At each stage the Infomap method included only links up to a certain length. The initial structure of 3 larger modules, formed by shorter range (< 2 mm) links is shown at bottom; intermediate structure (links < 6 mm) at middle, where nodes are now colored by the final module membership; and final structure with 8 modules, formed by all links, shown at top. Vertical lines show splitting of early modules into their ultimate components.

**S10 Figure.**
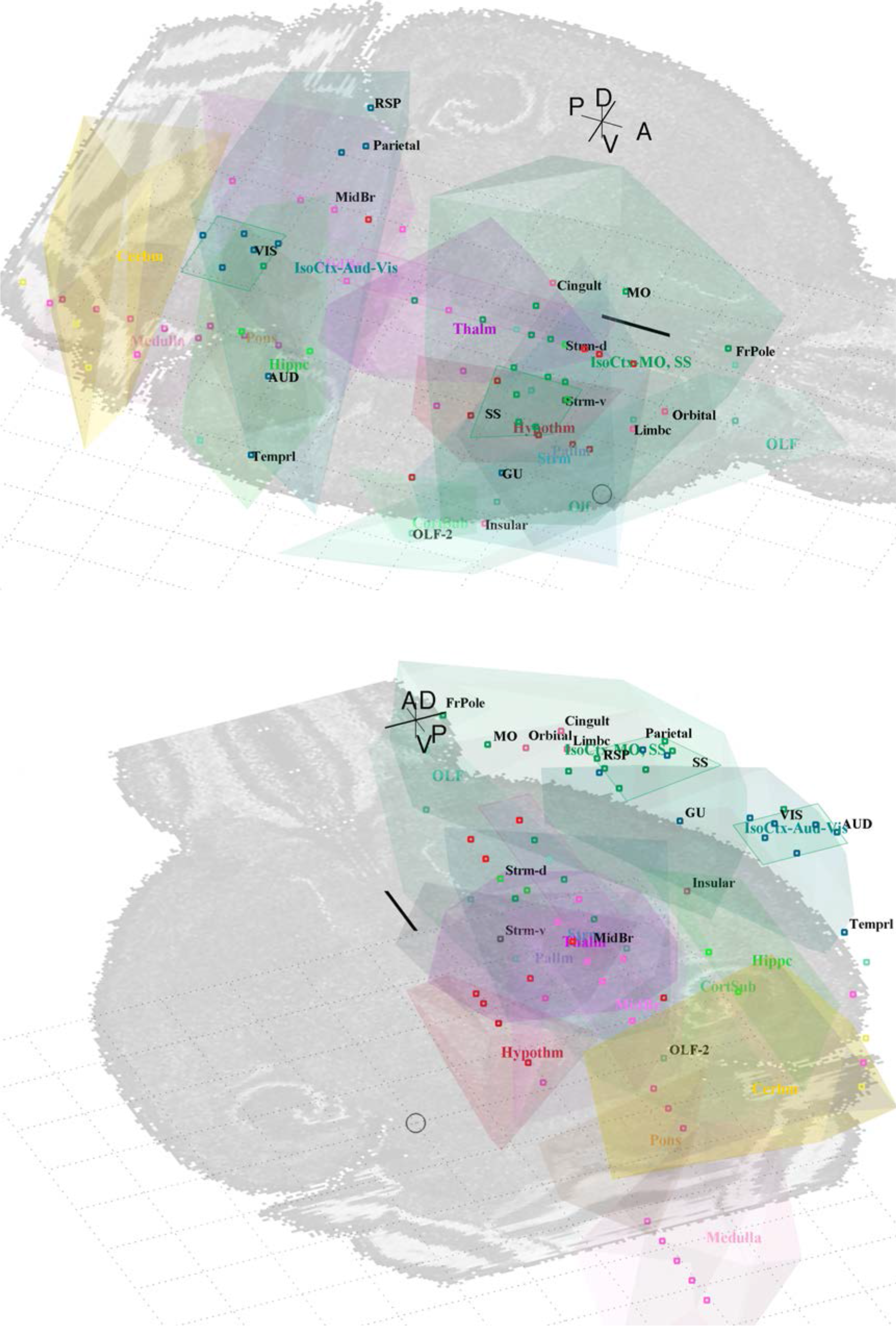
All 213 nodes plotted at the 45 schematic coordinates (listed in S3 data). The brain regions are represented by the encompassing convex hull (Fig 2), with other features as in S4 Figure. **a**. 3D perspective view from above the right front; **b**. view from left rear.

**S1 Table.** Top 20 nodes in order of decreasing probability flow calculated by the Infomap method [59–60], with other network measures listed. (59 KB PDF text file).

| ID | Acn | Region | Flow | Deg In,<br>wt | Deg Out,<br>wt | Deg In,<br>unwt | Deg Out,<br>unwt | rwC | nBC | Module |
| --- | --- | --- | --- | --- | --- | --- | --- | --- | --- | --- |
| 36 | CP | Striatum | 0.0588 | 453075 | 2360 | 87 | 22 | 0.0431 | 78 | 2 |
| 100 | MRN | Midbrain | 0.0486 | 182960 | 5647 | 88 | 77 | 0.0495 | 9 | 1 |
| 117 | PAG | Midbrain | 0.0330 | 164629 | 6368 | 98 | 101 | 0.0339 | 0 | 1 |
| 161 | SCm | Midbrain | 0.0314 | 215384 | 7160 | 78 | 74 | 0.0294 | 271 | 1 |
| 75 | LHA | Hypothalamus | 0.0288 | 149643 | 10634 | 110 | 114 | 0.0265 | 371 | 3 |
| 139 | PRNc | Pons | 0.0247 | 44790 | 6850 | 74 | 76 | 0.0278 | 106 | 1 |
| 168 | SNr | Midbrain | 0.0232 | 21911 | 8541 | 84 | 45 | 0.0217 | 512 | 2 |
| 57 | GRN | Medulla | 0.0207 | 33058 | 4130 | 58 | 47 | 0.0234 | 295 | 1 |
| 164 | SI | Pallidum | 0.0189 | 107414 | 3793 | 98 | 51 | 0.0164 | 0 | 3 |
| 140 | PRNr | Pons | 0.0173 | 65264 | 1685 | 84 | 24 | 0.0189 | 22 | 1 |
| 4 | ACB | Striatum | 0.0168 | 190736 | 7020 | 67 | 69 | 0.0127 | 0 | 3 |
| 68 | IRN | Medulla | 0.0166 | 42432 | 2775 | 59 | 35 | 0.0191 | 111 | 1 |
| 47 | ENTl | Hippocampal Formation | 0.0160 | 118185 | 34168 | 84 | 99 | 0.0124 | 431 | 4 |
| 55 | GPe | Pallidum | 0.0155 | 43915 | 6520 | 77 | 51 | 0.0135 | 53 | 2 |
| 129 | PIR | Olfactory Areas | 0.0145 | 112799 | 22312 | 93 | 45 | 0.0103 | 143 | 4 |
| 95 | MOp | Isocortex | 0.0137 | 105223 | 27542 | 79 | 119 | 0.0101 | 715 | 2 |
| 15 | AON | Olfactory Areas | 0.0130 | 38855 | 20315 | 81 | 41 | 0.0081 | 372 | 4 |
| 25 | CA1 | Hippocampal Formation | 0.0128 | 78005 | 2415 | 65 | 32 | 0.0101 | 105 | 6 |
| 94 | MOB | Olfactory Areas | 0.0125 | 45012 | 390 | 86 | 22 | 0.0081 | 1093 | 4 |
| 96 | MOs | Isocortex | 0.0124 | 119506 | 23453 | 75 | 133 | 0.0090 | 508 | 2 |

**S2 Table.** Likely hub nodes have high participation coefficient, P and weighted in-Degree or Out-Degree (30 KB PDF text file).

| ID | Acron | Region | P | Deg In, wt | Deg Out, wt |
| --- | --- | --- | --- | --- | --- |
| 31 | CLA | Cortical Subplate | 0.7860 | 19,666 | 117,022 |
| 47 | ENTl | Hippocampal Formation | 0.7877 | 118,185 | 34,168 |
| 32 | CLI | Midbrain | 0.7128 | 177,177 | 17,7177 |
| 135 | PP | Thalamus | 0.6987 | 2,775 | 152,884 |
| 75 | LHA | Hypothalamus | 0.6839 | 149,643 | 10,634 |
| 143 | PT | Thalamus | 0.6697 | 104,753 | 104,753 |
| 96 | MOs | Isocortex | 0.6450 | 119,506 | 23,453 |
| 89 | MGd | Thalamus | 0.5871 | 3,769 | 112,833 |
| 4 | ACB | Striatum | 0.5750 | 190,736 | 7,020 |
| 117 | PAG | Midbrain | 0.5204 | 164,629 | 6,368 |
| 184 | STN | Hypothalamus | 0.4923 | 6,624 | 171,909 |
| 129 | PIR | Olfactory Area | 0.3823 | 112,799 | 22,312 |
| 95 | MOp | Isocortex | 0.3439 | 105,223 | 27,542 |
| 161 | SCm | Midbrain | 0.3210 | 215,384 | 7,160 |

**S3 Table.** Comparison of Infomap modules with those found by the Louvain and the Newman spectral bisection methods. Infomap modules are list in approximately Anterior-Posterior and sensory-motor-midbrain-hindbrain order (24 KB PDF text file).

| Infomap Module | Function / Region | Louvain Module | Newman spectral |
| --- | --- | --- | --- |
| 4 | Olfactory | 1 | 1 |
| 2 | Somatosensory / Motor | 4 | 2 |
| 5 | Vis, Aud, TEa, RSP, PTLp | 6: Aud, TEa 5: Vis, RSP, PTLp | 2 |
| 7 | Association cortical areas | 2, 4, 5 | 2 |
| 3 | Hypothalamus | 1 | 1 |
| 6 | Hippocampus | 3 | 1 |
| 1 | Midbrain, Pons, Medulla | 1 | 1 |
| 8 | Cerebrum | 1 | 1 |

**S1 Text.** Additional details and discussion relevant to Methods and Results (154 KB PDF text file).

## Acknowledgments

The author thanks P Robinson and P Gong for support and valuable discussions, and acknowledges B Fulcher for valuable suggestions.

## Author Contributions

Conceived and designed the experiments: BAP. Performed the experiments: BAP. Analyzed the data: BAP. Contributed reagents/materials/analysis tools: BAP. Wrote the paper: BAP.

