## Supplementary material for "Network analysis of mesoscale mouse brain structural connectome yields modular structure that aligns with anatomical regions and sensory pathways": MouseBrain_S1text

### S1 Text.

Additional details and discussion relevant to Methods and Results are presented below.

#### Methods

#### Connectivity data.

The original AIBS dataset has been described elsewhere [9, 16, 18]. The fluorescent detected in target anatomical regions provides a measure of the strength of the directed links between those source and target regions. The connection strength was defined in four ways [9, 16, 18], starting from the total volume of fluorescent signal detected in 100 mm voxels across a target region, normalised variously by the injection and/or target site volumes. Those volumes varied in size from 70 to 24,000 voxels, with the majority less than 5,000 voxels (i.e. 5 $mm^3$).

#### Definition of connection strength.

The original raw data is the normalized connection strength, originally termed $w_{XY}$ [9], being the volume of fluorescent voxels detected in the target region arising from the volume of tracer injected in the source region. Here I discuss whether this is the true network connection strength between the two regions. That is a question of volume sampling in the source region: a) did the injection volume sample all relevant axons projecting to the chosen target region? And b) did the viral injection infect a sufficient fraction of the source region to diffuse to all linked targets within the timescale of the experiments (up to 21 days [9])? For relatively larger regions, with fewer outward links and smaller injection volumes it is possible to miss some efferent axons. That effect can be estimated by calculating a sampling metric for each region and experiment (injection volumes are listed in Supp. Table 2 of [9]). The metric $S_i = (Vol_i/k_i^{out})/\Delta V_i^{INJ}$, where $Vol_i$ is the source region volume, $k_i^{out}$ is the number of Out-links (or unweighted degree) from the source region (cf. S2b Figure) and $\Delta V_i^{INJ}$ is the mean injection volume. This calculation assumes the following: That the source region projects to $k_i^{out}$ separate targets each of which, on average, accepts source axons from a fractional volume ($Vol_i/k_i^{out}$) of the source region; And that the injected volume $\Delta V_i^{INJ}$ does sample that fractional volume and so sense the relevant outgoing axon or fibre bundle. The use of multiple injections, as reported [9], increases the chance of correctly sampling relevant portions of the source region. S2b Figure shows that most regions have a large number of out links, with the most probable number being ~40-50, with mean 60. Reported injections had an average volume of 0.24 $mm^3$, in a range of 0.002 – 1.36 $mm^3$. While some regions are quite large (eg. CP, MOB at 24 and 17 $mm^3$), the mean volume is 1.8 $mm^3$ while 56% are < 1 $mm^3$ and only 20 regions are > 5 $mm^3$. The latter contain likely candidates for under sampling. For $S_i < 1$ out links are likely to be well sampled since injected volumes, and their number, are large enough to span a sufficient volume of the source region. For $S_i > 1$ the injected volume is too small to sample all out links. The calculated metric was $S_i < 1$ for 96% of the 213 regions analysed, indicating that their out links are likely to be well sampled. For many of the smaller regions the injected volume spilled over into neighboring regions, necessitating a linear decomposition of signals to multiple separate target regions [9]. The calculated metric was $S_i > 1$ for nine cases, indicated those regions' Out-links were under sampled: FL, MDRNd, PAR, CP, MOB, Vispl, CA1 and PFL. Some (FL) were small with few out links, while others (CP, MOB) were large (24 & 17 $mm^3$) regions. All had 32 or fewer out links. The third largest region (PIR, 13 $mm^3$) had a marginal average score, $S_i = 1.18$, but the largest of those 7 injections (0.74 $mm^3$) yielded $S_i = 0.39$, possibly indicating good coverage. Of the 20 largest regions 5 were under sampled. By contrast CA3 is a relatively large region (5 $mm^3$) with few out links (32), so was flagged, but was well sampled (9 injections: 0.04 – 0.54 $mm^3$) so $S_i = 0.85$.

The extent of AAV infection of the source and all targets can be estimated with knowledge of the diffusion coefficient of AAV, measured as $D = 1.4\ \mu m/s^2$ [77]. For smaller injections of larger source regions time must be allowed for the viral tracer to diffuse across the region, to likely sources for distal targets. Typically the largest distance to travel would be half the size of a source region, which are in the range 0.2-1.5 mm. Order of magnitude estimates using the classical diffusion relation, $\langle r^2 \rangle = 6Dt$, indicates a worse case of order 10 days for the AAV tracer to diffuse over such distances. Subsequent transport along axonal pathways to target regions has also been modelled [78]. Overall the experimental timescale (up to 21 days [9]) appears to suffice for complete diffusion across source regions and along axons to targets. These tests indicate that the majority of measured normalized projection strengths are likely to reflect the true weight of source-target links, with the exceptions identified above. Thus use of the raw weights (cf. Methods) is justified, aside from the nine flagged cases where links weights are underestimated.

#### Data thresholds and normalisation

The raw connectivity, or adjacency, matrix normalised as described above contained 16,954 entries. These include 90 connections that returned to the source region. Thus there are 16,864 directed links between the 213 separate regions: 38% of all possible links. The 90 self-loops would constitute diagonal elements in the adjacency matrix. In network terms these self loops are links that are traced from a source region/node back to itself. Given that some regions have large voxel counts it is to be expected that sub-regions may connect between themselves. However, at the resolution of mesoscale connectivity, no further information is available. Theory shows that these self-loops can be significant in functional correlation matrices [15]. Most network analysis ignores such "return to source" links, and they are ignored in the present analysis. Numerical values of the adjacency matrix are presented in S1 Data.

The initial analysis [9] utilises an effective cut-off strength of $10^{-3.5}$ ($mm^3$), or approximately 1/3 of a voxel volume. This neglects the weaker 2762 structural links. That cut-off was also used to re-scale the link weights. Thus the weakest remaining link was assigned weight 1, so then the strongest link has weight 64.5k. A simple plot of all 16,854 detected links is revealing. A log-linear plot of the original experimentally measured link strength vs. link distance (ie. link length) is shown in S1 Figure, being the original quantitative connection strength (Suppl. Table 3 of [9]), that was described as the amount of segmented signal detected in the target region by infecting the source region [9]. S1 Figure highlights points lying beyond the main cluster of all other data, and that are likely to be outliers. The remaining cloud of data points shows more coherence and a clear trend with link distance.

It is widely recognised that the weakest measurements may constitute noise or be influenced by artefacts. Further some of the detected structural connections may not develop into functional links. Thus there is the question of choosing a threshold for the weakest links, and of establishing the correlation with functional links. The measured structural connectivity strengths span a large dynamic range with the weakest measurements being $10^{-16}$ below the strongest. Such a range might be commonplace in astrophysics but appears unusually large in the present context. The weakest ten measurements alone are 2-6 orders of magnitude below the main group of data and stand out as outliers in S1 Figure. The challenges of thresholding functional linkage measurements have been discussed in the context of generating binary, or unweighted, connectivity matrices [79]. Likewise the choice of threshold for the present structural data bears further study. The present structural linkage data, with its large range of weights, is amenable to examination by combining statistical data analysis methods with visualization. A standard approach is to plot and then de-trend the data, examine the distribution of residuals, and choose criteria for rejecting outliers, most simply taken as data lying beyond three standard deviations from the (local) mean. Given the trend in the weight – distance log-linear plot it is clear that applying a simple threshold penalises longer range links, with proportionally more discarded. This can then inform the choice of threshold, which may need to be linkage distance dependent. Possible roles of the weak links bears further investigation (cf. Results). Recent study showed that the links across large distances can have a role in global synchronization, based on a coupled oscillator model [80]. Choosing only the top 40% of structural links, consistent with ROC (cf. Methods) analysis [9, 45-46] discards a majority of the structural data. It does however reduce the risk of including non-functional links and aids the analysis and visualisation. The significance and possible roles of the weaker putative links is examined separately in the main text.

There is clearly a large variation in the raw connectivity data, suggesting possible sensitivity of the analysis to outliers in the data. Some care is required in applying the usual type of simple statistical tests, such as rejecting data more that three standard deviations from the mean, since that may eliminate potentially important links. Additional care needs to be taken to identify the correct sigma for the log-transformed data [81]. Beyond that, statistical measures of outlyingness [82], data depth [83] and data concordance tests can inform such analysis. Similarly the physics and biology of the system can guide acceptance and rejection of outliers: important transitions in actual physical and social systems often exhibit data excursions of more than three standard deviations from the mean.

The original data [9] have been re-analysed and also compared to expert curation of a selected subset of original experiments [9, 18], to examine the effects of noise and co-injection thresholds [18]. Other studies have analysed the correspondence between structural links, measured by tracers, and connectivity measured by MRI diffusion tractography [45]. And others [46, 68] focussed on links in the mouse isocortex, or the visual cortex [86]. Statistical analysis of the tradeoffs between True- and False-Positives rates, represented in so-called Receiver Operating Characteristic (ROC) curves [85], provides a measure of the correspondence between different type of connectivity measurements. The MRI tractography comparison produced a true-positive rate of 61% and a false-positive rate of 26% [45], suggesting a quantifiable approach to thresholding. Using a family of ROC curves at varying thresholds indicated that selection of the top 40% of structural links improved the correlation between structural and functional (rs-fcMRI) measurements [46]. The correspondence between the original raw measurements and expert curation (cf. Extended Data Figure 7 of [9]), of a random sample of 20 images, indicated a False Positive rate of 14.5% at the threshold $a_0$ ($10^{-3.5}$ $mm^3$), mostly due to image artefacts or low signal levels, possibly confounded with noise. Thus the present study focuses on the top 40%, by weight, of structural links by applying a link weight threshold at weight 78 using the normalisation described above.

A model based argument can cast light on that scaling. Using the average density of $10^5$ neuron/$mm^3$ in mouse cortex [86] suggests that about 3 neurons would fit in the cutoff volume. Thus the minimum link included in the adjacency matrix is plausibly close to one axon at this resolution, as noted by others [18]. The 14,192 links now included have strengths varying by nearly $10^5$, as noted previously [9]. Each axon, on average, might have 100-1000 synapses [87-88], so plausibly links of weight $10^{-2}$ should also be examined in subsequent sensitivity analysis. Using known average densities of neurons and synapses in mouse cortex [86] it can be estimated that, on average, those source and target regions may contain between 6.5k and 2.2m neurons, with most having around 500k neurons.

#### Network analysis and modular decomposition

Standard network measures have been applied to brain data [1, 5, 48], including to the mouse data [9, 16], and are used to rank the importance of nodes and links. These measures reflect different features of the underlying network structure [61, 89] and probe different types of process occurring on the network. Centrality measures probe more global aspects of the network such as how non-adjacent nodes communicate and can identify which are the "central" nodes in the network. Node degree is the simplest such measure and reflects local structure – that is nearest neighbour links, or paths of length one step. Degree also yields the long-time probability of visits by a random walker to the node [29, 33]. For directed links in-degree and out-degree are distinguished. Node strength, or weighted degree, generalises the concept of degree to a weighted network. Other measures are the node betweenness centrality (nBC) [49] and edge BC (eBC) [6, 28]. These commonly used centrality measures are based on graph or network concepts such as shortest graph paths or geodesics. Those count the number of intervening topological steps between nodes. This is distinct from the geometric path length, that reflects the distance travelled between nodes along links in physical space.
From graph theory the Dijkstra algorithm [49] facilitates computation of the shortest paths and is embodied in many of the standard codes [52-54]. Note that the un-weighted

links are used for that calculation. The restrictions described can be problematic, so care is needed in applying the methods and interpreting the results. For instance, nodes with many low-weight links may be emphasised over those with few high weight links. The standard methods do not take into account the spatial locations of neurons and the spatial embedding of the network structure; nor do they allow for routing of signal flows via indirect pathways as well as via shortest paths. They are appropriate to phenomena that embody targeted flow such as package delivery via a specified route in a transportation network, or disembodied interactions such as in a social network, rather than spreading phenomena on networks, such as diffusive flows or random walks. In brains where signals or pulse trains, that may encode information transmitted over axons and synapses, can spread by both direct paths and also via secondary links or more circuitous paths. This can have both deterministic and stochastic elements, so a random walk centrality measure (rwC), developed in other contexts [33], may be relevant. Node degree and node centrality measures (nBC, eBC, rwBC) were calculated by standard methods and toolboxes using available Matlab codes [52-54].

#### Temporal networks: signal delays

Time dependent phenomena can unfold on and spread over the fixed network [35-37]. The signalling is not instantaneous on the network; rather signals (both individual action potentials and pulse trains) travel along the network of axons at a fixed velocity. Thus both the network structure and the biophysics of signalling on the network determine where and when signals arrive at waypoints and putative targets. Signal velocity may vary between axons since it depends on axon diameter and degree of myelination [90]. Measurements of signal delays in a mouse thalamus-M1-ALM (pre-motor) circuit [9] indicate an axon conduction velocity of order ~ 1m/s, or equivalently 1 mm/millisec (mm/ms). This ignores variability with axon diameters and myelination. Raman scattering microscopy of mouse brain indicates ~10% region to region variation of myelin density [90]; the images indicate fairly constant myelin thickness along fibre tracts. Absent further information it is assumed here that all signal velocities over links are constant. The analysis also needs estimates of synaptic delay for multi-hop signal paths. A representative synaptic delay of 2 ms, measured in mouse neocortical pyramidal neurons [91], and signalling speed (i.e. axon velocity) of 1 mm/ms, adds an equivalent path length of 2 mm for the purposes of visualizating 2-step links. Thus link length (mm) is taken as a proxy for signal delay (in ms) over that link.

#### Schematic coordinates and 3D visualization

Plots of even medium sized networks with many links quickly become too dense to be intelligible. While many such visualizations are presented as network layouts in 2D, most can be generalised to 3D graphics, the latter requiring zooming, panning and rotations to identify and follow linkages. The brain is intrinsically a 3D structure, where spatial proximity is important for signalling so we need methods that enable its visualization in 3D. Neuroscientists often use 2D projections, sometimes in orthogonal pairs, to represent the anatomical layout brain regions.

A standard challenge in information visualization is choosing a simplified, schematic representation of a complex system, such as a network, that captures the essential features of the system and aids insight by presenting a 2D or 3D image that promotes intuition [92]. In so doing it is essential not to misrepresent structural or other important information. A long standing approach to presenting complex, spatially embedded networks is a schematic layout. Beck's 1932 schematic layout of the London underground rail network [93] is the classic example, variants of which have been widely adopted in major cities. It enables commuters to find their route in the complex network of rail links at a glance. Rather than faithfully tracing the rail links on the city map, a schematic map is developed. This simplifies major geographic features, such as the Thames river, to be nearly linear. The nodes (stations) are laid out on an almost rectangular grid. Linkages are generally straight lines drawn at a few regular angles. Together these steps reduce the visual congestion. In so doing it is important to keep a balance and not omit important structure. Overall the graphic captures the essential features of the network and omits myriad geographic details.

Noting the visual complexity of S4 Figure, it may be beneficial to test a simplified schematic layout of the mouse brain network. To that end, a set of schematic coordinates was developed, in which the original 213 anatomical regions, located at the Allen Atlas coordinates, were condensed onto 55 locations which each aggregated related and nearby regions (eg. combine ORBl, ORBm, ORBvl together; and ACAv, ACAd; and AUDd, AUDp, AUDv; and GU, VISC; and RSPd, RSPv, RSPagl; and ECTm TEa; for the mid- and hind-brain regions nodes were gathered into 2-4 groups along the anterior-posterior axis); the 35 sub regions of the Thalamus were divided into 6 groups, and the Isocortex into 14 groups. Typically these are closer to each other than the region's size (cf. voxel count in S2 Data). Vis and SSp are a special cases with each encompassing several separate sensory pathways, so their nodes were assembled into a small square grid rather than a single point. A guide was provided also by the essentially cylindrical symmetry of the anatomical regions in the Allen Atlas: with a clear anterior-posterior progression, and also an approximately radial separation of medial vs. lateral regions in a single hemisphere. Regions were typically divided into 2-5 sub-regions; however the isocortex, with more identified functional regions, was sub-divided into 22 sub-regions. The chosen coordinates are listed in S3 Data, where they are listed by region and in order of anterior-posterior location, and illustrated in S10 Figure. A benefit of this approach is that related links, often between nearby related nodes, become bundled into a few pathways, thereby enabling navigation of the 3D visualizations. They can be un-bundled, as required, to uncover local linkage details. Now all nodes and all links can be plotted in a graph with some prospect of navigating the link pathways. Subsequent figures place nodes at these schematic coordinates. Similarly, the 3D convex hull surfaces (cf. Figure 2) are used to represent brain regions is a schematic approximation. Not being space filling it causes an "exploded view" in 3D of brain regions and reduces crowding in the images. Surface colours match those presented in the Allen Brain Atlas sagittal slice images [63]. All figures were produced in Matlab, with all codes available [64].

#### Results

Additional details below describe results presented in Figures 4-6 and S5-S8 Figures.

#### Primary sensory signalling pathways

Cross modal linkages are evident in primary sensory pathways (Figure 4a). Both Visual and Auditory outputs have close – i.e. fast acting – and strong cross linkages: Vis-l to Aud-d & -p (d < 2mm, weights 1185, 630 respectively), Vis-am to Aud-d (d < 3mm, weight 150); and Aud-d to Vis-p (d < 2mm, weight 845), Aud-v to Vis-al (d < 2.5mm, weight 790) and Aud-p to Vis-am (d < 4mm, weight 330). Both Auditory and Visual outputs also return to regions in the Midbrain (to Ic-e, -d, SCm and to SC-s, -m respectively). GU/Visc regions link to SS-s and the Insular, Temporal and Parietal regions, and also, less strongly, to the Hypothalamus (LHA); GU alone also links to SSp-bfd, -m while Visc alone links to SSp-bfd, -m, to SSs, to MOs and to the olfactory region PIR. Olf, multiple Somatosensory and GU/Visc links converge to CP, along with Auditory inputs. Visual inputs are not direct, but rather via Parietal and RSP regions. Somatosensory and GU/Visc regions link to the posterior association areas, with the latter also interacting with the ventral Olfactory region (PIR); also note that SS links to the Retrosplenial region, while GU/Visc links to Parietal. The GU/Visc regions link directly to: Auditory (GU to Aud-p, -v), to Somatosensory (GU to SSp-m; Visc to SSp-tr, -ll, SSs), Visual (GU to Vis-p, -am; Visc to Vis-p, -l, -pm, -pl) regions, and to the Cortical Subplate (Claustrum). At this long distance the Frontal Pole only receives strong input from the Orbital association area. Overall most interactions are within module.

Figure 4 and S5 Figure highlight 5 network hubs, as discussed in the main text. The next hub-like target is the endopiriform nucleus (EPd, Cortical Subplate) with medium strength (weight > 1.5k) In-links from the Cortical Subplate (CLA, BLA), Isocortex (AIv, PERI), Olf (PAA, TT) and Midbrain (CLI). The caudoputamen (CP, Striatum) is the largest sampled region [9] and likely in consequence has a large number (72, cf. S2b Figure) of strong In-links. The strongest link in the connectome is from the Frontal Pole to CP (weight 64k). Other very strong links (weight > 10k) are from the Cortical Subplate (BLA), the Isocortex (PERI, ACAd), Thalamus (AMd, PF, CM) and Midbrain (SNc, CLI). The next suggested hub is the dorsal peduncular region (DP, Olf) has only relatively weak (weight < 500) inputs, the strongest of which are from the Thalamus and PERI. Next is the postsubiculum (POST, Hippocampus) with modest (weight > 500) links in locally from SUBd, and then from Midbrain (NOT), the Retrosplenial region (RSPv), Thalamus (AD, AV) and Cortical Subplate (Claustrum). The perirhinal region (PERI, Isocortex) has medium (weight > 1k) links in from the Cortical Subplate (LA, CLA), Olf (PAA) and Thalamus (SMT, SPFm), along with a weaker (weight 556) input from the frontal pole.

#### Association Areas

Figure 5 displays In- and Out-links of the anterior cortical association areas (Limbic, Orbital, Insular and Cingulate). In total there are 354 strong (top 40%by weight) In-links (Figure 5a) to these four regions; sensory inputs are listed first. Orbital regions receive sensory inputs: ORB-l from AON, weight 190; ORB-l, -vl from SSp-ll, weights 90, 125; ORB-vl from SSp-bfd, -tr, weights 135, 115; ORB-vl from VISC, weight 545. Longer range inputs are: ORB-l, -vl, -m from AUD-v, weights 330, 265, 115; ORB-vl from Vis-l, weight 415. Limbic regions receive inputs: ILA from AON, weight, 175; PL from SSp-ll, weight 105; PL and ILA from VISC, weights 140, 245; PL and ILA from Vis-l, weights 80, 340; ILA from AUD-v, weight 245. Insular regions receive inputs: AI-d, -p from GU, weights 1.2k, 300; AI-v, -p from SSp-n, both weights 95; AIp from SSp-m, weight 110; AId from AUD-v, weight 220; AId from VISal, weight 295. Cingulate regions receive inputs: ACA-d, -v from SSp-ll, weights 425, 85; ACAd from SSp-tr, weight 255; ACAd from SSp-bfd, weight 85; ACAv from GU and VISC, weights 135, 770; ACAv from VISp, weight 215; both ACA-d, -v from VIS-l, weight, 295, 335; ACAd from AUDv, weight 90. AId has a very strong input (weight 13k) from the Frontal Pole, with weaker intputs from AIv (weight 1.2k), AIp (weight 678) and the prelimbic region (weight 1.4k) to FRP. Overall these association areas share the inputs from the more anterior senses, and also have lesser inputs from the posterior senses: AUD and VIS. There are also inputs from the posterior association areas: ORB-l, -vl from PTLp, weights 200, 480; ACA-d, -v from PTLp, weights 865, 460; ILA and PL from PTLp, weights 250, 145; ILA from RSPv, weight 140; ORBl from TEa, weight, 80; AId from TEa, weight 275. All these anterior regions receive strong (weights ~1k - 2k) structural links from the Claustrum, as found by other tracer studies of the mouse neocortex [68].

Figure 5b displays the 399 strong (i.e. top 40% by weight) Out-links from the four anterior cortical association areas. (AIv to MOs, weight 6.6k; AId to MOp, weight 1.1k; ORBl and ORBm to MOs, weights 4.5k and 1.3k, respectively; ORBvl to MOp, weight 1.1k; ACAd to MOs, weight 4.8k; ACAv to MOp, weight 500). Only ORB-l, -m connects to Frontal Pole (weights 635, 200). Feedback to sensory regions is: from Insular to Olfactory; from Limbic, Orbital and Insular to Somatosnsory regions (ILA to SSp-tr, weight 120; ORBvl to SSp-m, weight 80; ORB-l, -vl to SSp-ul, weights 100, 360; ORB-vl to SSp-ll, weight 170; ORB-l, -vl to SSp-bfd, weights 540, 480; ORB-vl to SSp-tr, weight 340; AIp to SSp-n, -ul, weights 200, 90; AIv to SSp-tr, weight 80; ACAd to SSp-tr, weight 660); while groups of three regions feedback to Auditory and Visual regions (ILA to Vis-p, weight 300; ORB-vl, -l toVis-p, weights 420, 100; AId to Vis-al, weight 620; ACAd to Vis-pm, -p, -l, -pl, weights 100, 2.2k, 270, 90; and ILA to Aud-d, weight 170; ORB-l to Aud-v, weight 80; AIv to Aud-v, weight 190; ACAd to Aud-d, weight 200). There are medium strength outputs to the Frontal Pole, from: ORB-l, -m, weights 635, 205; and AI-v, -d, -p, weights 1.2k, 345, 120. Longer range links are to the posterior cortical association areas: ORBvl, ACAd, ACAv to RSPagl, weights 160, 360, 520; AIp to RSPv, weight 155; AIp to RSPd, weight 85; ORBvl, ACAd to PTLp, weights 695, 3.5k; ILA, ORBvl, AIv, ACAd to TEa, weights 160, 330, 1.5k, 200; ORBm, AIv, ACAd to ECT, weights 170, 1.8k, 360). All these anterior regions have medium strength (weights ~100-400) structural links to the Claustrum; the Out-Links are significantly weaker than the In-Links from Claustrum.

Figure 6a displays the strong inputs to the three posterior association areas (Temporal, Parietal and Retrosplenial). The nearby auditory-visual regions provide immediate strong inputs: AUD-v, -d to TEa, weights 2.5k, 2.3k; AUD-d to ECT, weight 700; AUDp to PTLp, weight 805; AUD-d, -p to PTLp, weights 3.6k, 805; VIS-l to ECT, weight 2.5k; VIS-p to TEa and ECT , weights 80, 255; VIS-am to TEa, weight 630; VIS-am to ECT, weight 575; VIS-l to RSP-agl, -d, -v weight 1.1k, 2k, 550; VIS-pl to RSP-v, weight 550. Other sensory inputs are: VISC to TEa and ECT, weights, 260, 275; VISC to PTLp, weight 3k; VISC to RSPv, weight 1.5k; VISC to RSP-agl, -d, weights 930, 1.9k; SSp-bfd to TEa and ECT, weights 405, 225; SSp-tr to TEa, weight 90; SSp-ll to both TEa, ECT, weights 235, 85; SSp-bfd, -tr, -ll to PTLp, weights 310, 730, 305. Weaker sensory inputs are AUD-v to PTLp, weight 140; VIS-am to RSP-d, -v, weight 135, 140; VIS-p to RSP-agl, -v, -d, weights 290, 215, 405; VIS-pl to RSP-d, weight 190; VIS-l to RSP-d, weight 190; GU to RSPv, weight 125; SSP-tr to RSPd, weight 85. The Temporal regions receive multiple inputs from the Hippocampus (SUBv, ENTl, ENTm, PRE, PAR, POST, SUBd). Pairs of the posterior regions receive inputs from all four anterior regions: Orbital, Insular and Cingulate to RSP and Temporal; and Limbic to PTL and Temporal regions. The Motor regions input to RSP and Temporal. There is very strong Thalmic input: MGd to TEa, weight 20.8k; and AD to RSPv, weight 31k. There are weaker, long range inputs from the Frontal Pole: to ECT, weight 295; and to TEa, weight 685; along with feedback from MOs to both ECT and TEa, weights 205, 200. Again the anterior regions receive strong structural links from the Claustrum.

Figure 6b displays the immediate feedback from posterior cortical association areas to the local sensory regions: PTLp to AUD-d –p, -v, weights 1.2k, 1.3k, 220; PTLp to VIS-l, -p, -pl, -am, -pm weights 660, 1.9k, 130, 1.8k, 580; PTLp to SSp-tr, weight 1.3k; PTLp to VISC, weight 1.8k; RSPagl to Vis-pm, weight 1k; RSPv to VIS-am, -pm, -p, weights 630, 220, 1k; RSPagl, -d to VIS-pl, weights 120, 95; RSP-d, –v to SSp-tr, weights 740, 1k; RSP–v to SSp-ll, weight 130; RSP–v to SSp-bfd, weight 270; RSP–v, -agl, -d to VISC, weights 560, 235, 690. Both RSP-d and PTLp link to Midbrain regions: RSP-d and PTLp to SCm, weights 2.3k, 2.8k; RSP-d and PTLp to PAG, weights 945, 475; PTLp also links to numerous other Midbrain regions: SCm, SCs, Ice, ICc, APN, MRN, NPL and SNr. The three regions all have relatively weak (weights ~100-200) links to numerous Thalmic regions: notably strong links are: ECT to LD, weight 810; PTLp to LP, weight 835; RSPv to LD, weight 1.3k. RSPagl links to the Hippocampal fields CA1, CA3 with weights 175, 120. There are direct links to Motor regions: TEa to MOp and MOs, weights 760, 425; PTLp to MOp and MOs, weights 1.1k, 2k; RSP-d, -v to MOp, weights 860, 1.1k. The links to the anterior association areas are: TEa to ORB-l, weight 80; PTLp to ACA-v, -d, weights 460, 865; PTLp to ORB-l, -vl, weights 200, 480. The retrosplenial regions only have structural links to the Cingulate regions: RSPd to ACAd, weight 940 and RSPv to ACAv, weight 1.8k.

Links with the Frontal Pole are displayed in S7 Figure and discussed in the main text. Other strong inputs to FRP (cf. S7a Figure) are from Claustrum, Thalamus and Olfactory (NLOT) regions. There is a weaker long range input from Aud-v. The very strong outputs from FRP are to the Insular regions (weights 13k and 27k) and Motor region, MOs (weight 22k). The strongest link in the mouse connectome appears here: FRP to Caudoputamen (weight 64k). Other medium strength outputs are to SSp-m, -ul, -n and SSs (weights 2k, 3.8k, 300, 2.5k); along with Vis-al (weight 550). There are weaker, long range links to the Temporal association areas (ECT, TEa, weights 680, 300) and the Perirhinal region (weight 550). Other output are to ventral regions: Claustrum, Striatum, Palladium, Hippocampus and Midbrain. there are only a few strong inputs to FRP, from: ORB-l, weight 635 and AIv, weight 1.2k; and Thalamus (VM, weight 600). The other 14 inputs considered are all medium strength (weights ~100, 200). Weak links are presented in S7b Figure. Also evident in the image is a longer range link to the Retrosplenial region, and a number of 2-step links to RSP and the Parietal region. There are noticeable, medium weight, feedback links into FRP from anterior sensory regions: Olfactory (AON) and Somatosensory (SSp-m ,-ul, -bfd, SSs). Other weak inputs are from: Palladium, Hypothalamus, Hippocampus, Midbrain, Pons and Medulla.

#### Motor regions

S8 Figure show the strong links with MOp and MOs. Strong inputs are from the Cingulate (ACAd to MOs, weight 4.8k; ACAv to MOp, weight 500), Orbital (ORBl to MOs, weight 4.5k; ORB-vl, -l to MOp, weights 1.1k, 1k), Insular (AI-v, -p to MOs, weights 6.6k, 800; AI-d, -p to MOp, weights 1.1k, 945) and Retrosplenial and Parietal regions (RSP-d and PTLp to MOs, weights 2.8k, 2k; RSP-v, -d to MOp, weights 1.1k, 880). Temporal input is from TEa to MOs and MOp, weights 425, 760. Direct sensory inputs are from multiple Somatosensory primary regions (SSp-ll, -tr, -ul, -bfd to MOs, weights 1.1k, 990, 845, 2k; SSp-tr, -ul, -bfd, -m, -n to MOp, weights 1.4k, 3.7k, 2.7k, 4k, 1.2k; and SSs to both MOs and MOp, weights 790, 1.6k), along with Gustatory and Visceral links to MOs (weights 1.7k, 1.4k, respectively) and to MOp (weights 4.9k, 930). There are a strong inputs from Aud-v (weights 2k, 1.4k) and Vis-am (430, 300) to both MOs and MOp; and relatively weak inputs from AUD-d to MOs only, weight 385; and AUD-p and VIS-al to to MOp only, (weights 320, 335). Olfactory input is: DP to MOp, weight 305. Note that other direct inputs from the Olfactory regions are relatively weak; rather AON links to motor regions via Orbital and Insular association areas. Hippocampal input is CA2 to MOp, weight 125.

Nearby outputs are numerous medium strength links to thalmic areas: MOp links to MD, PO, PF, RT, VPL, CM, VAL, VPMpc, and VM (listed in order of increasing distance; weights ~ 130 – 300), with a weaker (weight 30) link to CL; MOs has feedback to Vis-am and Aud-p & -v, and again outputs to CL, MD, PO, RT, PF, VAL and VM. MOs also has feedback to associations areas: ACAd, ORB-vl & -l, AI-d and –v, PERI, ETC and TEa. MOp has strong feedback to AId (weights 350) and weaker structural links to ACAv, FRP, ORB-l and PTLp (weights 5, 50, 10, 20). Both MOp and MOs have longer range links to Striatum, Midbrain and most hind brain regions. Tracer experiments [94] have investigated the spatial organisation of L5 and L6 outputs from MOp and MOs, finding a segregation into a core of MOp projections to thalamic areas, enveloped in a shell of MOs projections.
